# Data-Constrained Recurrent Network Neural Model Uncovers the Circuit Mechanism of Olfactory OFF Responses

**DOI:** 10.64898/2026.05.22.727331

**Authors:** Shruti Joshi, Autumn K. McLane-Svoboda, M. Gabriela Navas-Zuloaga, Camron Stout, Debajit Saha, Maksim Bazhenov

## Abstract

Sensory neural circuits must encode both the presence and termination of a stimulus. Following odor offset, projection neurons (PNs) in the insect antennal lobe (AL) exhibit transient increases in firing rate, termed OFF responses, yet the circuit mechanisms that generate them in recurrent excitatory-inhibitory networks remain poorly understood. Here, we constructed a biologically-constrained firing rate-based recurrent neural network (RNN) model of the locust AL and trained it on electrophysiological recordings from 110 in vivo PNs to reconstruct their odor-evoked temporal dynamics across five odorants. The trained model faithfully reproduced the firing rates of constrained neurons, while unconstrained PNs and LNs developed biologically plausible temporal response patterns and response-type diversity. Using targeted input and connectivity perturbations, we found that OFF responses arise through two mechanistically distinct pathways. A feedforward pathway transmits offset-type olfactory receptor neuron (ORN) input directly to downstream PNs, while a recurrent pathway generates post-stimulus excitation independently of offset input. Selective perturbation of individual recurrent connections identified LN-LN mutual inhibition as the dominant recurrent pathway, and decomposition of excitatory and inhibitory inputs revealed that it produces net excitation through the transient release of inhibition rather than through increased drive. The two pathways recruit largely non-overlapping PN populations, indicating that OFF response identity is an emergent property of the network state rather than a cell-intrinsic feature. These findings provide a circuit-level account of OFF response generation and demonstrate how data-constrained RNNs can dissect circuit mechanisms directly from *in vivo* recordings.

## Introduction

Sensory neural circuits must encode not only the presence of a stimulus but also its termination. In olfaction, the cessation of an odor is a behaviorally significant event as it signals the end of a stimulus encounter, resets ongoing sensory representations, and may itself carry information about odor identity [Mazor et al., 2005, Saha et al., 2017]. At the circuit level, this is reflected in transient increases in neural firing rate that occur following stimulus termination (OFF responses), which have been observed across sensory modalities, including audition [Bondanelli et al., 2021, Scholl et al., 2010] and olfaction [Kim et al., 2023, Nagel and Wilson, 2016, Saha et al., 2017]. Despite their ubiquity, the circuit mechanisms that generate OFF responses in recurrent excitatory-inhibitory networks remain poorly understood.

The insect antennal lobe (AL) offers a tractable system for investigating these mechanisms. As the first central relay of the olfactory pathway, the AL receives input from olfactory receptor neurons (ORNs) in the antennae and transforms it through a dense recurrent network of excitatory projection neurons (PNs) and inhibitory local interneurons (LNs) [Laurent, 2002]. PNs relay processed odor representations to higher brain areas, while LNs sculpt network activity through lateral and recurrent inhibition [Laurent, 2002, Laurent et al., 1996, Saha et al., 2013]. In the locust, odor stimulation evokes complex spatiotemporal patterns of PN activity that unfold over the course of seconds, with distinct neural ensembles activated during stimulus presentation (ON responses) and following its termination (OFF responses), and some responses that show increased firing at both ON and OFF time periods (ON-OFF responses) [Laurent et al., 1996, Mazor and Laurent, 2005, Mazor et al., 2005, Saha et al., 2013, 2017]. ON and OFF ensembles are largely non-overlapping and encode orthogonal representations of stimulus presence and absence [Saha et al., 2017].

Two candidate mechanisms could generate OFF responses in the AL. The first is a feedforward mechanism: a subset of locust ORNs exhibit offset-type temporal response motifs, firing preferentially at stimulus termination [Kim et al., 2023], and could directly drive post-stimulus excitation in downstream PNs [Scholl et al., 2010]. The second is a recurrent mechanism: the dense inhibitory connectivity within the AL, particularly among LNs, could generate post-stimulus excitation through network-level dynamics independent of the temporal structure of the sensory input [Nagel and Wilson, 2016, Saha et al., 2017]. Intracellular recordings from the locust AL have provided evidence consistent with the latter possibility, showing that disengagement of recurrent inhibition coincides with increased PN firing at odor offset [Saha et al., 2017]. However, isolating the specific circuit motif responsible and determining how feedforward and recurrent contributions interact has remained challenging using experimental approaches alone, as selectively manipulating individual synaptic pathways while preserving the rest of the circuit is difficult *in vivo*.

Computational modeling provides a principled framework for addressing these questions. Existing models of the locust AL have employed hand-tuned spiking or firing-rate networks to investigate oscillatory synchronization, odor discrimination, and learning-dependent modulation of odor representations [Bazhenov et al., 2001, 2005, 2013, Chen et al., 2015, Joshi et al., 2025, Patel et al., 2009]. While these models have provided valuable insights into AL function, manual parameter adjustment introduces experimenter bias, limits the model’s capacity to discover unanticipated solutions, and does not fully exploit the high-dimensional neural data now available from modern recording techniques. Recurrent neural networks (RNNs) trained via gradient-based optimization offer a powerful alternative. Task-trained RNNs have revealed interpretable dynamical motifs underlying context-dependent computation, sequential activity, working memory, and flexible multitask processing [Driscoll et al., 2024, Dubreuil et al., 2022, Kim and Sejnowski, 2021, Mante et al., 2013, Rajan et al., 2016, Sussillo and Barak, 2013, Sussillo et al., 2015, Yamins et al., 2014]. More recently, data-constrained RNNs — models trained directly on neural recordings rather than behavioral tasks — have been used to infer directed interactions between brain areas and uncover latent dynamical structure inaccessible from recorded activity alone [Durstewitz et al., 2022, Koppe et al., 2019, Perich et al., 2020]. Because these models are optimized under biological constraints, their learned parameters and dynamics can be interrogated through targeted perturbations to generate testable hypotheses about the mechanism, an approach that has proven particularly effective for linking circuit structure to neural dynamics [Durstewitz et al., 2022, Koppe et al., 2019, Perich et al., 2020, Valente et al., 2022].

In this study, we apply this data-constrained RNN framework to the locust AL to investigate the mechanistic basis of OFF responses in PNs. We construct a biologically-constrained firing rate-based RNN model [Kim et al., 2019] comprising 830 excitatory PNs and 300 inhibitory LNs, matching the known cell-type composition of the locust AL [Laurent, 2002], and train it on electrophysiological recordings from 110 *in vivo* PNs to reconstruct their odor-evoked temporal dynamics across five odorants. The optimized model faithfully reproduces the firing rate profiles of constrained neurons, while unconstrained PNs and LNs develop biologically plausible temporal response patterns and response-type diversity without direct training. We use this model to dissect the mechanisms underlying OFF response generation through a series of input and connectivity perturbations. By systematically removing ORN offset inputs, we demonstrate that recurrent network dynamics are sufficient to generate OFF responses, albeit in reduced numbers. Selective perturbation of individual recurrent pathways identifies LN-LN mutual inhibition and resulting disinhibition in PNs as the dominant recurrent mechanism, and decomposition of excitatory and inhibitory inputs reveals two distinct mechanisms for producing post-stimulus excitation. These findings provide a concrete, circuit-level account of how the AL generates OFF responses through parallel feedforward and recurrent mechanisms operating on distinct neuronal populations, and generate specific predictions for experimental validation.

## Results

### *In vivo* antennal lobe recordings to train RNN

Electrophysiology recordings were obtained from the antennal lobe (AL) of locusts using a multielectrode array positioned to capture activity from PN populations. During experiments, a standardized panel of five odorants: hexanal, hexenyl acetate, pentanal, propylbenzene, and mineral oil, mixed at a concentration of 1% vol/vol, were used during experiments to record odor-evoked response. Odor stimulus was delivered through a controlled olfactometer system (Fig 1A) lasting four seconds and repeated for five trials to assess response consistency across trials. Recording positions within the AL were chosen based on visual confirmation of odor-evoked responses using the voltage traces.

**Fig 1.**
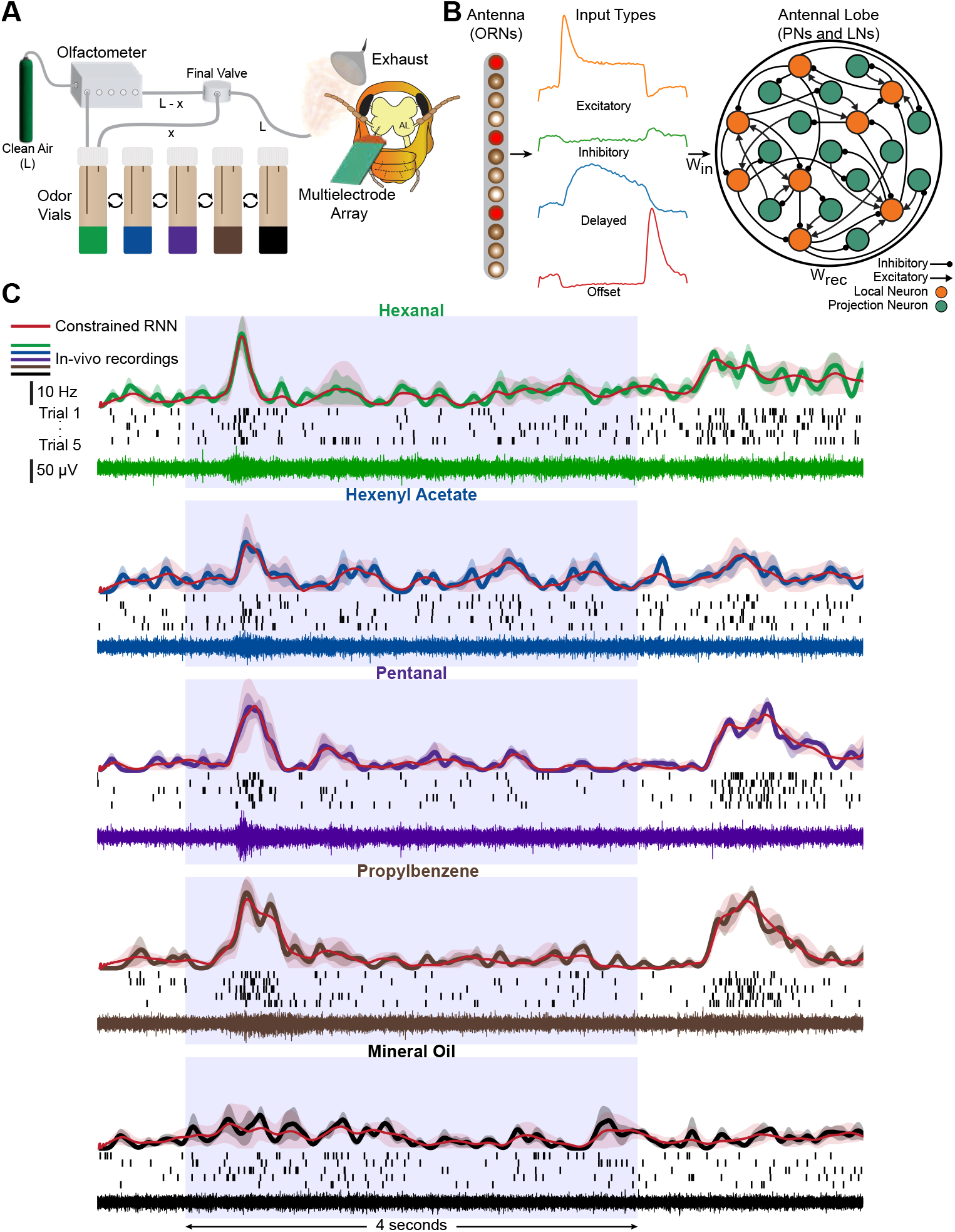
RNN activity closely replicates *in vivo* PNs. A) Schematic of experimental data collection setup showing olfactometer delivery system and multielectrode array recording from the locust antennal lobe (AL). Odor vials were attached to the olfactometer to control the timing of odor release. The entire flow rate (L) before stimulus is delivered to the locust’s antennae. During stimulus, a portion of the air (x) diverts into the odor vial headspace. The clean air (L-x) and the odor-laden air (x) are combined at the final valve and delivered to the antennae. A multielectrode array placed within the (AL) records changes in projection neuron (PN) activity from the odor-evoked responses. Each odor was presented five times. B) Architecture of the RNN model. Input connectivity from ORNs exhibits diverse temporal response profiles (excitatory, inhibitory, delayed, and offset motifs) to the AL network containing excitatory PNs and inhibitory LNs with recurrent connectivity (*W _rec*) and input weights (*W _in*). C) Comparison of *in vivo* PN firing rates (colored traces) with model-reconstructed firing rates (red traces overlaid on top of the colored traces) across the five odorants for a representative PN. One voltage trace (bottom) out of the five trials is shown. Spike-sorted voltage traces are displayed as raster plots (middle) across five trials, where each black line represents a spiking event. Individual peri-stimulus time histograms (PSTHs, top) derived from the raster plots illustrate changes in spiking rate through time with shaded regions indicate SEM across trials. Blue boxes indicate odor stimulus window (0 to 4 seconds).

Odor-evoked voltage traces (Fig 1C, bottom) displayed multi-unit activity from surrounding neurons with visual differences between odors. Individual PNs (*n* = 110 neurons) were identified by spike sorting (see Methods) the voltage traces to produce raster plots. All five trials were averaged to create peri-stimulus time histograms (PSTHs) of individual neuron firing rates. A representative neuron’s odor-evoked response is shown in Fig 1C, displaying a unique change in firing rate at the onset of odor presentation and after the odor is stopped (offset). When looking at raster plots of every neuron and trial (Fig S1), small variations in odor-evoked responses can be seen at the individual neuron level. While somewhat different, most neurons displayed an increase in spiking within the first second of odor stimulus (0 to 1 seconds) and the two seconds after stimulus ends (4 to 6 seconds). This response timing can also be observed when gathering neurons into population PSTH data, separated by odor (Fig S2C). At a population level, differences between odor-evoked responses become clearer, with firing rates for odors such as hexanal and pentanal being higher than those for hexenyl acetate and propylbenzene during stimulus onset. Mineral oil served as an experimental control, providing the RNN model with sufficient real-world examples of stimuli that elicited very little or no neural response when presented. During training, this allowed for robustness within the model, preventing the network from becoming overly sensitive while distinguishing between important response features and background noise. RNN construction was based on the makeup of the locust AL and trained using individual trial responses of each *in vivo* neuron with the aim of capturing the dynamics of odor-evoked responses.

### Continuous firing rate based RNN model of the antennal lobe

We constructed a biologically-inspired firing rate-based RNN model to simulate the computational dynamics of the locust AL [Kim et al., 2019]. The model architecture was designed to incorporate the two primary neuron classes in the AL circuitry: excitatory PNs, which serve as output units that relay olfactory information to higher brain centers, and inhibitory local interneurons (LNs), which provide lateral inhibition and sculpt network-wide response patterns through recurrent connectivity. To ensure biological realism, the model comprised 830 PNs and 300 LNs, directly matching the cell population sizes reported in anatomical studies of the locust AL [Laurent, 2002]. The temporal dynamics of each model neuron were governed by a first-order differential equation:

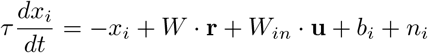

where *x*_*i*_ represents the synaptic drive to neuron *i, τ* is the effective integration time constant, *W* is the recurrent connectivity, **r** is the vector of firing rates of all the neurons in the recurrent network, *W*_*in*_ is the input weight connectivity, **u** is the vector of ORN inputs, *b*_*i*_ is the background synaptic drive to neuron *i*, and *n*_*i*_ is the random noise added to the activity. The instantaneous firing rate was computed as:

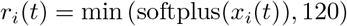

where the softplus nonlinearity provides a smooth rectification and the saturation at 120 Hz reflects reasonable constraints on maximum firing rates observed in locust AL neurons.

Both PN and LN populations received inputs from a population of 1000 simulated olfactory receptor neurons (ORNs) that provided the sensory drive to the network (Fig S2A). The ORN population was designed to exhibit the full spectrum of temporal response motifs observed in biological olfactory sensory neurons [Kim et al., 2023], including excitatory onset responses, delayed onset responses, inhibited responses, and offset responses. These diverse input patterns were generated using a computational model we previously designed that reproduced experimentally observed ORN spike train statistics and temporal dynamics [Kim et al., 2023], ensuring that the AL model received biologically plausible sensory input patterns (Fig 1B).

The network was trained using backpropagation through time (BPTT) to reconstruct the experimentally recorded firing rate profiles of 110 *in vivo* PNs. To achieve this objective, we implemented a training approach where 110 of the 830 model PNs were designated as “constrained PNs” whose firing rates were explicitly trained to match the temporal dynamics of their corresponding *in vivo* counterparts. The remaining 720 PNs, along with all 300 LNs, were designated as “unconstrained neurons” whose activities evolved freely according to the network dynamics, subject to biologically motivated regularization constraints that prevented unrealistic parameter values. The loss function was defined using root mean squared error (rMSE) between model and *in vivo* PN firing rates, providing a direct measure of reconstruction fidelity across the entire temporal response profile for each constrained PN. An additional regularization term in the loss function penalized the mean firing rate of a random subset of unconstrained PNs and LNs to match that of the recorded *in vivo* PNs, preventing unrealistic activity (see Methods). The parameters of the trained model are detailed in Fig S3.

### Trained AL RNN neurons reconstruct firing rates of *in vivo* PNs

After training by *in vivo* data, AL model reliably reconstructed the firing rate dynamics of *in vivo* PNs. Across all odors, constrained PNs generated firing rate profiles that closely matched experimental recordings in onset timing, activity magnitude and duration, and offset responses following stimulus termination (Fig S2D,E). Mineral oil, serving as a control odorant, consistently evoked weaker responses in both experimental and model neurons, confirming the specificity of odor-evoked activity patterns. Averaged across all constrained PNs and training trials, the model achieved a correlation of 0.72 ± 0.15 (mean ± SD) with corresponding *in vivo* firing rate profiles, with representative examples shown in Fig 1C.

Notably, unconstrained PNs also exhibited biologically plausible temporal response patterns consistent with *in vivo* observations, as reflected in the population-averaged activity across all 830 model PNs (Fig S2F,G). This indicates that recurrent dynamics shaped during training generalized beyond the constrained units, producing realistic temporal patterns across the entire PN population. Together, these results demonstrate the model’s effectiveness as a tool for investigating the computational principles underlying olfactory processing in the locust AL.

### Temporal properties of PNs in vivo and in the AL model

Recorded *in vivo* PNs exhibited a rich diversity of temporal response patterns to odor stimulation, including transient increases in firing rate at stimulus onset, sustained activity during odor presentation, and offset responses following stimulus termination, consistent with previous characterizations of AL dynamics [Saha et al., 2017]. To systematically characterize this diversity, we classified PN activity into four distinct response types: increased firing rate at stimulus onset only (onset or ON), increased firing rate following odor termination only (offset or OFF), increased firing rate at both stimulus onset and offset (onset-offset or ON-OFF), and unchanged firing throughout the trial (non-responsive or NR). Responses were classified based on an activity threshold calculated from baseline and a minimum firing rate threshold (see Methods for details). This classification was applied to both *in vivo* and RNN neurons, yielding 2,750 classified responses for *in vivo* data (110 neurons × 5 odors × 5 trials) and 20,750 for the entire RNN (830 neurons × 5 odors × 5 trials). Representative sorting examples from *in vivo* neuron trials can be seen in Fig S4.

As expected, *in vivo* PN response profiles varied across odorants (Fig 2C), consistent with previous studies showing that odor identity strongly influences the temporal structure produced by the AL [Laurent et al., 1996, Mazor and Laurent, 2005, Saha et al., 2013]. For hexanal, hexenyl acetate, and pentanal, onset responses were the most prevalent, with mean number of responses per trial ranging from 41.4–46.4 (out of 110 responses). Onset-offset responses were consistently fewer, while offset only responses were rare across these odors (mean numbers ranging between 7.8 to 10.6). Mineral oil, used as a control, showed elevated NR counts and substantially reduced onset-offset responses, consistent with the minimal neural activity reported above (Fig S2C).

**Fig 2.**
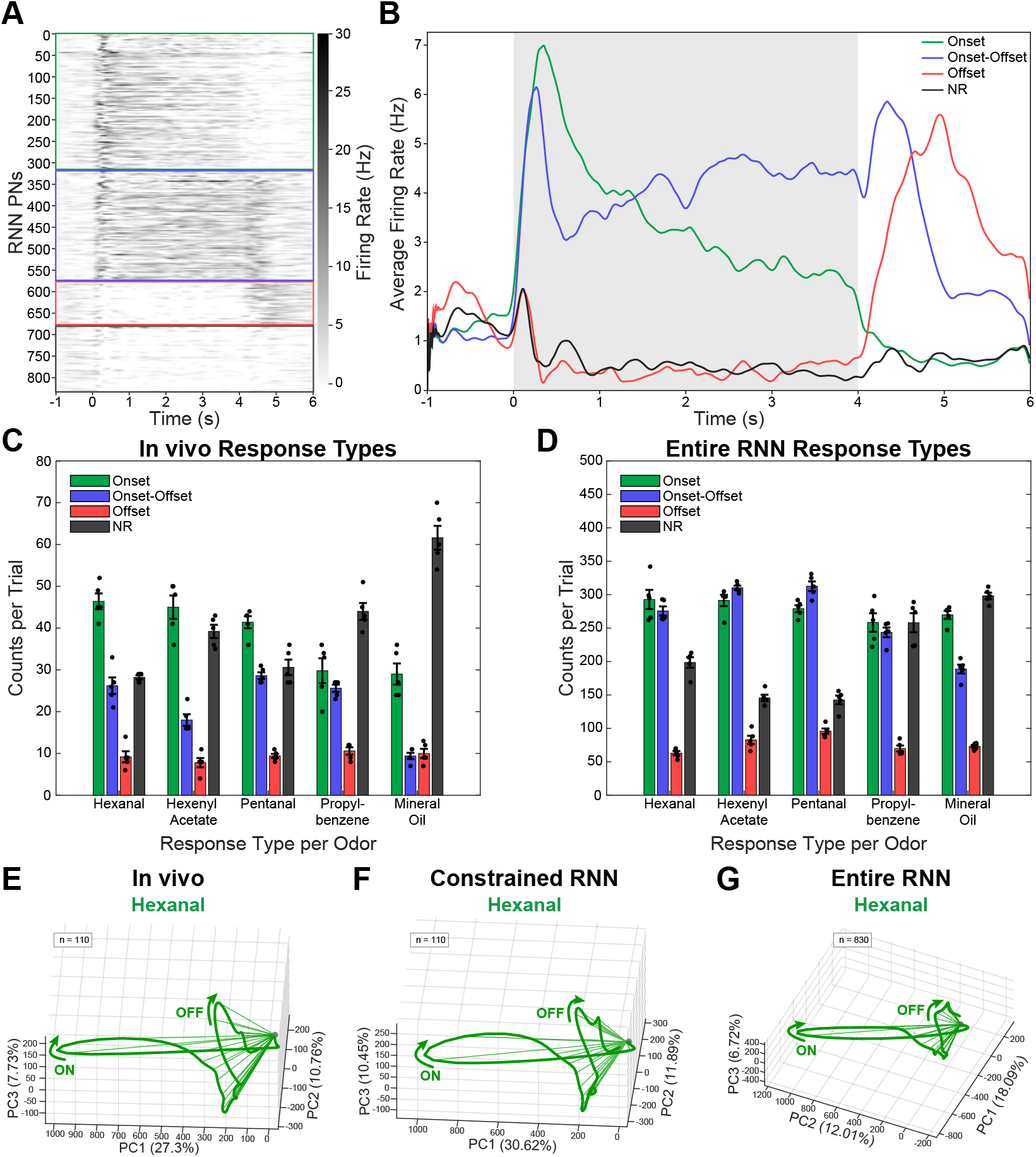
Temporal properties of *in vivo* and model PNs. A) Firing rate heat map of all 830 RNN PNs for one trial from a representative odor. Odor input is from 0 to 4 seconds. Green box depicts neurons that were classified as having an onset response to that specific odor, the red box surrounds neurons classified as an offset response, the blue box is onset-offset, and the black box is no response (see Methods for classification methods). B) Average firing rate profiles of onset (green), offset (red), onset-offset (blue), and no response (NR, black) type RNN PNs in response to stimulation (representative odor shown). C) Distribution of response types among experimentally recorded *in vivo* PNs (*n* = 110) across all five odorants. Most odors show more neurons exhibiting an onset response to odor stimulation, with fewer onset-offset and even fewer offset responses. Black circular points represent the number of neurons for that trial exhibiting a specific response to each odor; five points for each column due to having five trials. Error bars depict standard deviation between trials. See Fig S4 for individual representative classification examples. D) Distribution of response types in the optimized RNN model (*n* = 830) across the same odorant set. The model reproduces the predominance of onset and onset-offset type responses while generating all four classes. E) Principal component analysis (PCA) of temporal response dynamics for experimentally recorded *in vivo* target PNs (*n* = 110). Characteristic trajectory loops with distinct onset (labeled on) and offset (labeled OFF) response phases shown for hexanal. See Fig S6A for *in vivo* PCAs for other odors. F) PCA trajectories for constrained model PNs (*n* = 110), closely matching the temporal structure observed in experimental data with similar loop patterns and odor-specific trajectories. See Fig S6B for constrained model PCAs for other odors. G) PCA analysis of all model PNs (constrained and unconstrained, *n* = 830), demonstrating that the entire network population exhibits biologically plausible temporal dynamics similar, but not identical, to those observed in constrained PNs and *in vivo* recordings. See Fig S6C for entire model PCAs for other odors. -0.5 to 6 seconds is displayed in the PCAs, with odor onset at 0 and offset at 4 seconds.

The RNN model successfully reproduced this diversity of temporal dynamics (Fig 1C and S2D,E,F). Looking at the full population of raw data from one trial in Fig 2A, neurons sorted into the same response type (labeled with colored boxes) portray a homogeneous firing pattern, suggesting our sorting criteria is effective. For each response category, the average firing rate across all RNN PNs (Fig 2B) matches expectations: onset responses peak exclusively during odor presentation, offset peaking only after stimulus termination, and onset-offset during both periods. While the absolute proportions of response types differed from *in vivo* observations, the model recapitulated key trends: onset responses remained high across tested odorants (trial mean numbers range 258.4–292.8, out of 830 neurons), with fewer pure offset responses (mean numbers range 63–95.8). Mineral oil still produced the highest number of NR and the lowest number of onset-offset responses (Fig 2D).

Despite these differences in response type proportions, the trial-to-trial variability for neural activity was comparable between datasets, with similar median values and SD ratios close to 1 (Fig S5A,B), indicating that the model preserves realistic levels of variability rather than oversimplifying activity. Odor-specific trial-level variability was quantified as the standard deviation of firing rates across neurons within each trial. Bootstrap resampling over neurons revealed overlapping confidence intervals between *in vivo* and RNN neural activity after accounting for differences in neuron count (Fig S5A), suggesting that any observed differences are largely attributable to the RNN having more neurons while still maintaining biological plausibility.

To visualize onset and offset dynamics at the population level, odor-evoked neural trajectories were constructed using principal component analysis (PCA) to reduce dimensionality (see Methods). PCA was applied to each dataset for all odor inputs. The time window of interest was chosen from − 0.5 to 6 seconds, encompassing half a second of baseline, odor onset (at 0 seconds), odor offset (at 4 seconds), and a return to baseline. The first three principal components with the highest variance were chosen to project the data onto three dimensions. The odor-evoked vectors for every 50 ms time bin were connected temporally to show neural trajectories over time and aligned at the origin. PCA revealed that temporal response dynamics of constrained model PNs closely matched those of experimentally recorded neurons. Both populations exhibited characteristic trajectory patterns in PC space, with distinct “loops” corresponding to onset and offset response phases (Fig 2E,F and Fig S6A,B). Notably, the full model population, including unconstrained PNs, displayed similar temporal structure in PC space, indicating that the recurrent network dynamics produced biologically plausible response patterns across the entire PN population (Fig 2G and Fig S6C). These results demonstrate that the optimized AL model captures both the diversity of PN response types and the temporal dynamics observed in biological neural circuits.

### OFF responses persist in the absence of ORN offset input

OFF responses in the AL can arise through two distinct mechanisms in our model. The first is a feedforward mechanism in which PNs receive direct excitatory input from ORNs that exhibit offset-type activity, transmitting their temporal dynamics directly to downstream PNs. The second is a recurrent mechanism, in which the circuitry of the AL generates post-stimulus excitation through network-level dynamics that do not depend on the temporal structure of the feedforward input.

To dissect the contributions of these two mechanisms, we systematically perturbed the ORN offset input to the trained model while leaving all other ORN response types (excitatory, inhibitory, and delayed) intact. Specifically, we scaled the firing rates of ORNs exhibiting offset-type responses by factors ranging from 0.0x (complete removal) to 2.0x (doubling), and quantified the resulting changes in PN response profiles across the network. In all these experiments, the full intact model was trained initially by *in vivo* data as described above.

Even with complete removal of ORN offset input (0.0x perturbation), some PNs continued to generate OFF responses (Fig 3A,B). The average firing rate profile of these neurons exhibited a clear post-stimulus peak following odor termination (Fig 3B, red trace), confirming that these represent offset responses produced entirely through recurrent network dynamics. However, the number of pure OFF responses was substantially reduced compared to the unperturbed condition (Fig 3C, ∼ 20 responses per trial out of 830 total responses), indicating that while recurrent dynamics are sufficient to generate OFF responses, feedforward offset input is the dominant contributor under normal operating conditions.

**Fig 3.**
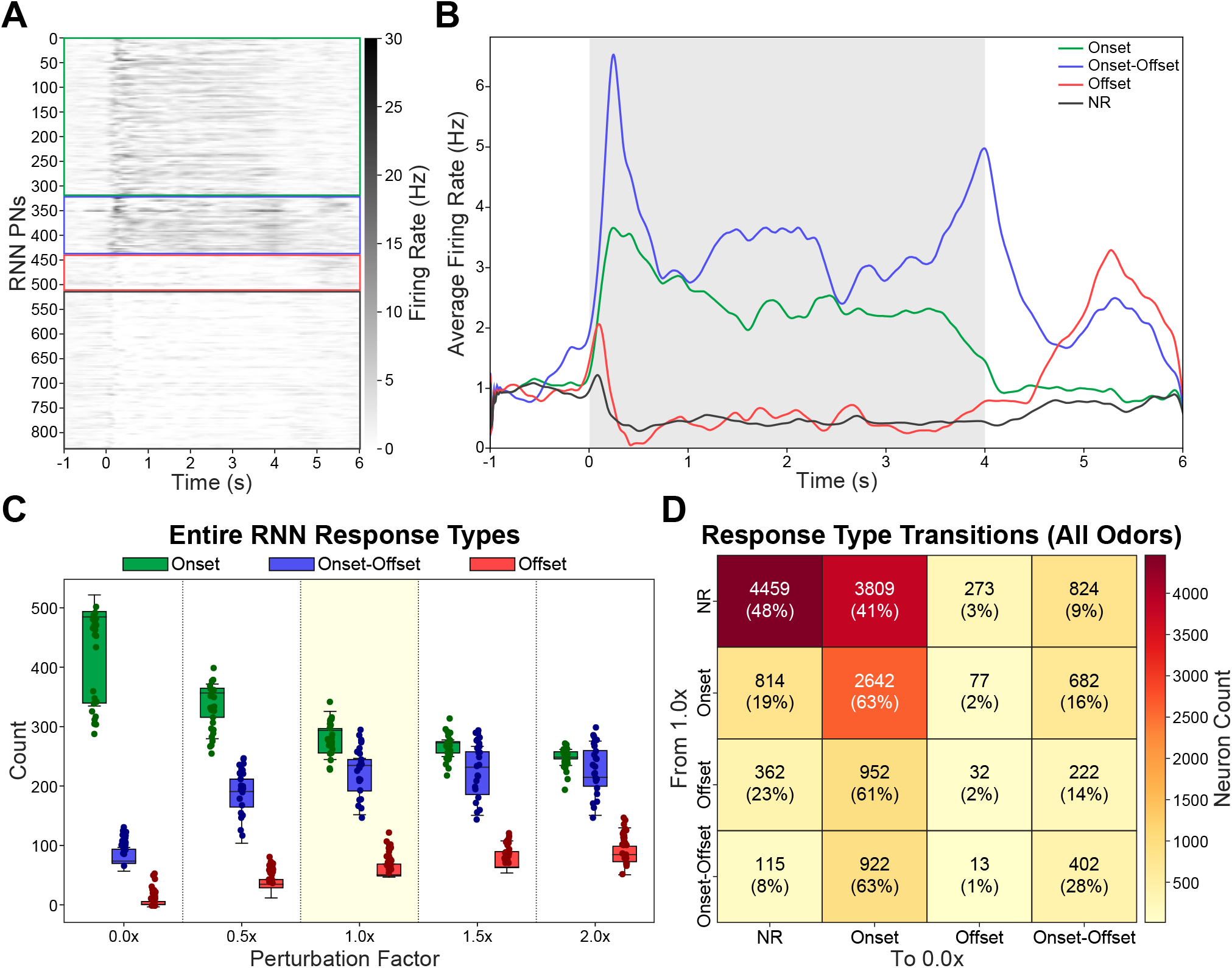
Effect of OFF-input perturbation on PN response types. A) Heatmap of PN firing rates in the absence of OFF inputs. A single trial for Pentanal is shown. PNs are sorted by response type, with boxes indicating boundaries between types. B) Average PN firing rate per response type in the absence of OFF inputs. Each line corresponds to one response type. The shaded area indicates odor presentation. [A single trial of Pentanal is shown]. C) PN response-type counts for different perturbation factors scaling connections form OFF inputs. [Response counts for each trial and odor are shown]. D) Transition matrix for PNs response type changes between 1.0x and 0.0x OFF input perturbation conditions across all odors.

To examine how the removal of offset input restructures response identity at the single-neuron level, we constructed a transition matrix tracking how individual PNs change their response classification between the 1.0x and 0.0x conditions across all odors (Fig 3D). This analysis revealed that removing offset input reorganizes functional identity across the entire network rather than simply eliminating a fixed subset of OFF responses. Of the PN responses classified as offset at 1.0x, the majority (61%) transitioned to onset responses at 0.0x while only 2% retained their offset identity, indicating that the offset-generating mechanisms recruit largely distinct neuronal populations between conditions. Similarly, 63% of onset-offset responses at 1.0x were reclassified as onset-only at 0.0x, consistent with selective loss of their offset component. Interestingly, a small subset of new offset responses emerged in the absence of the OFF inputs. This was likely a result of disinhibition. Because both PNs and LNs receive ORN input, the removal of ORN offset drive also eliminates offset-evoked LN activation, releasing specific PNs from inhibition and enabling offset responses in neurons that were previously suppressed during the offset epoch. The reorganization extended beyond offset-related neurons: 41% of non-responses (NR) at 1.0x emerged as onset responses at 0.0x, and 19% of onset responses became NR, demonstrating that ORN offset input shapes network dynamics throughout the entire trial rather than solely during the post-stimulus period. Together, these transitions indicate that OFF response identity is not a fixed, cell-intrinsic property but an emergent feature of the network state shaped by the interaction between feedforward drive and recurrent processing.

### LN-LN inhibition is the dominant recurrent pathway for OFF response generation

Having established that recurrent network dynamics contribute to the generation of OFF PN responses in the absence of direct ORN offset input (Fig 3), we next sought to identify which specific recurrent pathway is responsible. The AL circuit contains three connection motifs that could shape post-stimulus dynamics: LN-LN (mutual inhibition among local interneurons), LN-PN (direct inhibition of projection neurons), and PN-LN (excitatory drive onto local interneurons). To dissect the contribution of each pathway, we systematically scaled each set of connection weights by factors of 0.5x (reduced) and 1.5x (increased), with 1x being the intact model in the model with ORN offset inputs removed, and quantified the resulting changes in PN response type counts. As previously, all these experiments started from the pretrained intact model.

Scaling LN-LN weights produced the strongest effect on OFF responses among all pathway-specific perturbations (Fig 4D, second row). Reducing LN-LN inhibition to 0.5x nearly eliminated recurrently generated OFF responses, whereas increasing it to 1.5x enhanced OFF response counts beyond the intact condition. Onset-offset responses, which include a post-stimulus component, showed the same dependence on LN-LN strength (Fig 4C, second row), consistent with their offset phase also being generated by LN-LN inhibition. Reducing LN-LN weights also decreased onset response counts (Fig 4B, second row); however, this reduction was accompanied by a decrease in overall PN firing rates (Fig 4A, second row), suggesting it reflects a nonspecific drop in network excitability rather than a selective disruption of onset dynamics. To disentangle these effects, we performed a global inhibition perturbation in which both LN-LN and LN-PN weights were scaled together, thereby maintaining comparable average PN firing rates across perturbation levels (Fig 4A, top row). Under these conditions, reducing global inhibition to 0.5x mostly preserved onset response counts but still nearly abolished OFF responses (Fig 4B,D, top row). This dissociation confirms that the loss of OFF responses upon reducing LN-LN weights reflects a specific disruption of the recurrent mechanism generating them, rather than a general consequence of reduced network excitability.

**Fig 4.**
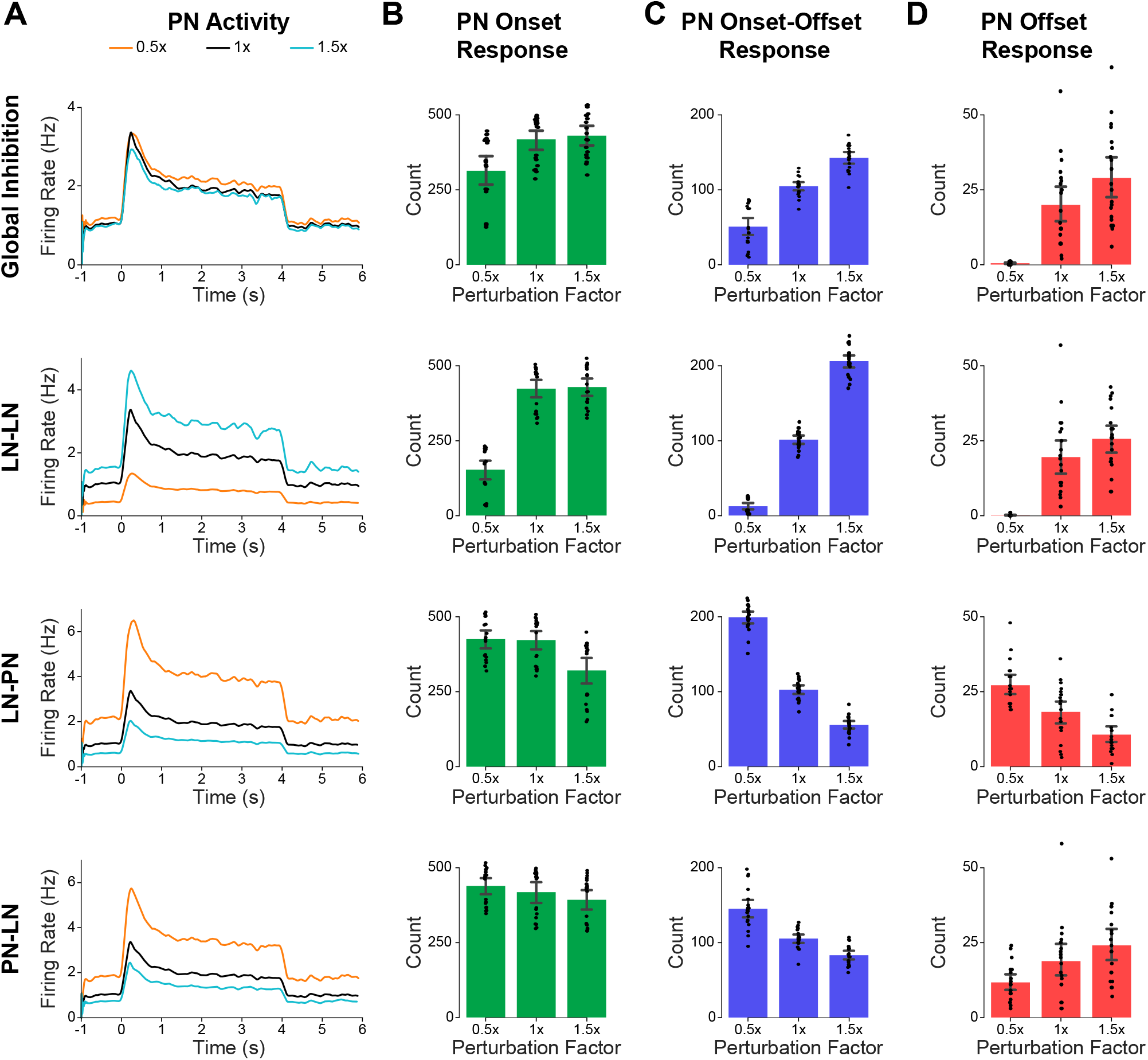
Effect of recurrent connectivity on PN responses in the absence of OFF inputs. A) PN average firing rate for the model without OFF inputs, with specific recurrent connection weights either reduced by 50% (0.5x), intact (1x), or increased by 50% (1.5x). From top to bottom, manipulated weights are: all inhibitory weights (“Global Inhibition”), only LN-LN, only LN-PN, or only PN-LN. B) Count of PN onset (ON) responses across all trials and odors for specific recurrent weights reduced by 50% (0.5x), intact (1x), or increased by 50% (1.5x). Rows are as in A. C) As B, for PN Onset-Offset (ON-OFF) response counts. D) As B, for PN Offset (OFF) response counts.

In contrast, scaling LN-PN weights alone primarily modulated overall network excitability rather than selectively affecting OFF responses. Reducing LN-PN inhibition to 0.5x elevated PN firing rates across the entire trial (Fig 4A, third row) and broadly increased response counts across all categories (Fig 4B-D, third row). PN-LN perturbation had the weakest overall effect on OFF response counts (Fig 4D, bottom row), consistent with this pathway serving as an excitatory feedback loop rather than directly mediating post-stimulus inhibitory dynamics.

Together, these results identify LN-LN mutual inhibition as the primary recurrent pathway underlying OFF response generation in the absence of feedforward offset input. The selective dependence of OFF responses on LN-LN connection strength points to a disinhibitory circuit motif as the recurrent source of post-stimulus PN excitation.

### Distinct excitatory-inhibitory dynamics underlie feedforward and recurrent OFF response mechanisms

Having identified LN-LN inhibition as the dominant recurrent pathway for OFF response generation (Fig 4), we next examined the synaptic mechanisms by which feedforward and recurrent pathways produce net excitation at stimulus offset. We decomposed the total input onto OFF-responding PNs into excitatory (E) and inhibitory (I) components, computed as deviation from pre-stimulus baseline (Δ input; see Analysis of RNN model responses), across three conditions.

In the intact model, OFF responses were driven by a feedforward mechanism: offset ORN input directly excites PNs while co-activating LNs, causing both E and I to rise at offset, with excitation slightly exceeding inhibition to yield a positive E-I difference (Fig 5A). In the absence of offset inputs, OFF responses were instead generated through disinhibition: excitatory input returned to baseline, but inhibitory input dropped below baseline (Fig 5B), producing net excitation through withdrawal of inhibition rather than through increased drive. Removing LN-LN connections on top of offset input removal abolished this disinhibitory dynamic; neither E nor I showed the transient changes needed to produce net excitation, and OFF responses were eliminated (Fig 5C). The signed area under curve (AUC) of E-I across all OFF PNs confirmed these patterns: the intact model showed a broad distribution with a large positive region, the offset-removed condition was narrower but still predominantly positive, and removing LN-LN connections shifted the distribution toward negative E-I AUC (Fig 5D).

**Fig 5.**
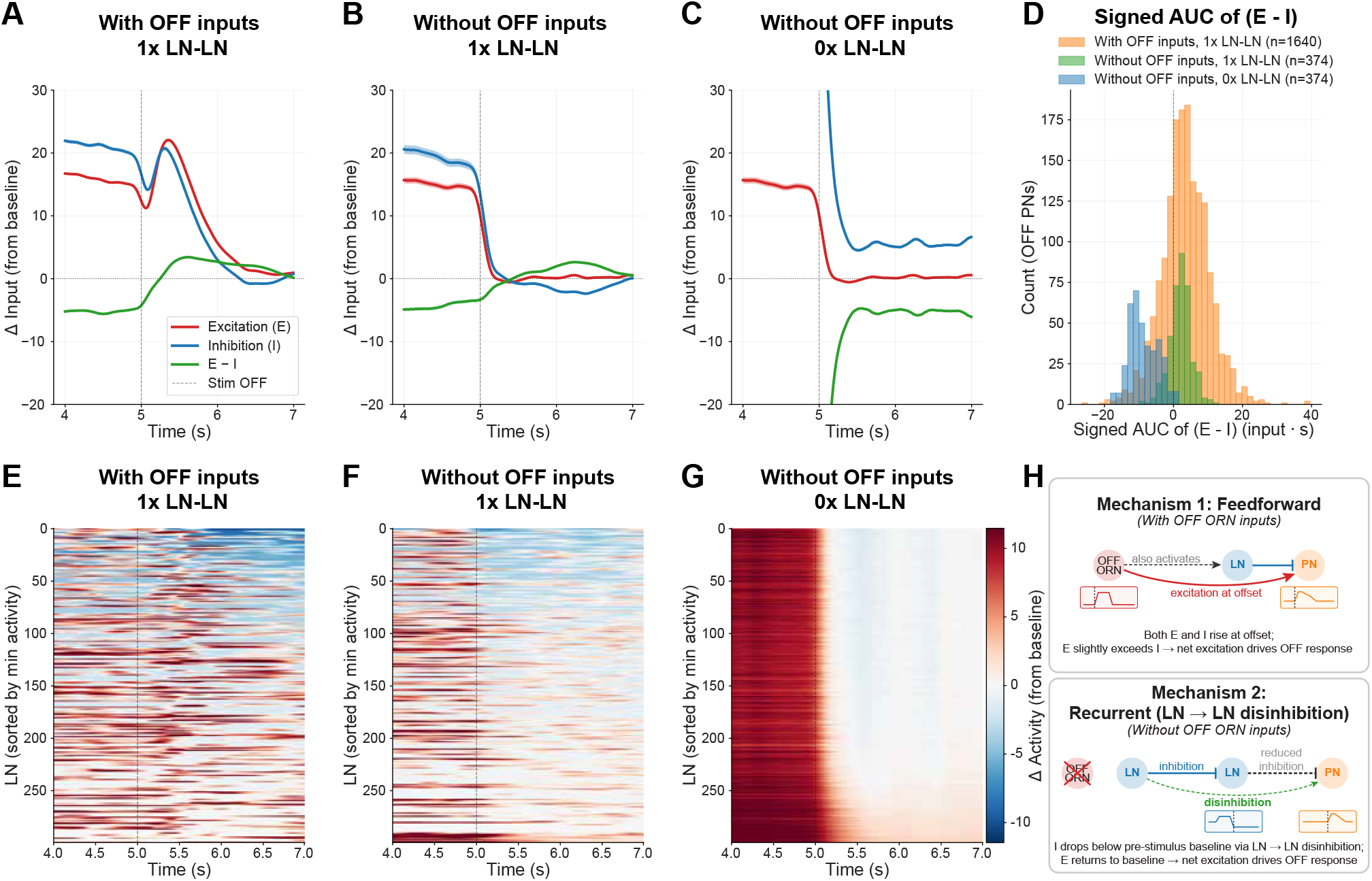
Mechanisms of PN OFF response generation. A) Average Excitatory (E) and Inhibitory (I) inputs to PNs with OFF responses across odors and trials in the original unmodified model. Inputs are computed as the product of pre-synaptic activities and synaptic weights, and are presented as difference from baseline (Δ input), where baseline is the average input before odor presentation. E-I denotes the difference between average excitatory and inhibitory inputs, with E-I *>* 0 after the end of odor presentation (“Stim Off”, vertical line) resulting in an OFF response. B) As A, for PN OFF responses generated in the absence of OFF inputs. C) As A, when both OFF inputs and LN-LN connections are removed. PN OFF responses are not generated in this condition; averages are obtained for the same PNs considered in B to illustrate how their inputs change. D) Distribution of the area under the excitation minus inhibition curve for three conditions shown in A, B, and C. Positive values on the X-axis indicates more excitation than inhibition and negative values show more inhibition than excitation over the entire offset time period. E) Heatmap of baseline subtracted firing rates for LNs, sorted by peak minimum activity. F) As E, for the model without OFF inputs. G) As E, for the model without OFF inputs or LN-LN connections. H) Schematic showing the two mechanisms for generation of OFF responses.

The mechanistic basis of the recurrent pathway was visible at the LN population level. In the intact model, LNs showed diverse activity at offset (Fig 5E). Without offset inputs, a prominent LN subpopulation was suppressed below baseline at offset (Fig 5F). This suppression reduces inhibitory drive onto downstream PNs, consistent with a disinhibitory mechanism, and recruits PNs that were not part of the OFF population in the intact model. Despite the absence of offset input, LN activity remained heterogeneous, indicating that LN–LN interactions sustain population-level diversity (Fig 5F). Removing LN–LN connections abolished this dynamic entirely, eliminating LN suppression at offset and preventing recurrently generated OFF responses (Fig 5G). Together, these results confirm that LN–LN mutual inhibition is required for the disinhibitory cascade underlying recurrent OFF response generation.

Thus, the AL employs two distinct mechanisms to generate OFF responses (Fig 5H): a feedforward pathway that drives post-stimulus excitation through direct ORN offset input, and a recurrent pathway that achieves the same outcome through an entirely different route, the transient release of inhibition via LN-LN mutual inhibition.

## Discussion

In this study, we constructed a biologically-constrained firing rate-based RNN model of the locust AL, trained on electrophysiological recordings from 110 *in vivo* PNs to reconstruct their odor-evoked temporal dynamics across five odorants. The trained model reliably captured the firing rates of constrained neurons, while unconstrained PNs and LNs developed biologically plausible temporal response patterns and response-type diversity (Fig 1, Fig 2). Unlike RNN training paradigms that prioritize task performance [Driscoll et al., 2024, Kim and Sejnowski, 2021, Lappalainen et al., 2024, Mante et al., 2013], our approach trains the network directly on *in vivo* recordings, allowing gradient-based optimization to determine connectivity weights, time constants, and biases within biologically motivated constraints. Using this framework, we identified two mechanistically distinct pathways for generating OFF responses, i.e., transient excitatory response observed after odor termination [Nagel and Wilson, 2016, Saha et al., 2017], in the AL: a feedforward pathway driven by ORN offset inputs to the AL and a recurrent pathway mediated by LN-LN disinhibition, each operating on largely non-overlapping neuronal populations (Figs 3, 4, 5).

Our work addresses a central challenge in computational neuroscience: bridging the gap between neural activity recordings and the circuit mechanisms that produce them. Existing computational models of the locust AL have relied on hand-tuned parameters to reproduce experimentally observed activity patterns [Bazhenov et al., 2001, 2005, 2013, Chen et al., 2015, Joshi et al., 2025, Patel et al., 2009], yielding valuable insights into contrast enhancement, sparsification of odor representations, and concentration-invariant coding. However, manual parameter adjustment introduces experimenter bias, limits the model’s capacity to discover unanticipated solutions, and does not fully leverage the high-dimensional neural data now available from modern recording techniques. Recurrent neural networks offer a powerful alternative [Barak, 2017], as they can learn directly from empirical data and capture complex temporal dependencies, making them well-suited for modeling the intricate dynamics of neural circuits. A growing body of work has begun to exploit this capacity, demonstrating that RNNs can not only reproduce neural activity but also yield mechanistic insights into the circuits they model. Task-trained RNNs have revealed how recurrent dynamics give rise to context-dependent computation, sequential activity, working memory, flexible multitask processing, and more, while identifying interpretable dynamical motifs such as line attractors, selection vectors, and low-rank population structure that illuminate the computational strategies employed by neural circuits [Driscoll et al., 2024, Dubreuil et al., 2022, Kim and Sejnowski, 2021, Lappalainen et al., 2024, Mante et al., 2013, Rajan et al., 2016, Yamins et al., 2014]. Building on this foundation, data-constrained RNNs trained directly on multi-region recordings have been used to infer directed interactions between brain areas and uncover latent dynamical structure that would be inaccessible from recorded activity alone [Durstewitz et al., 2022, Koppe et al., 2019, Perich et al., 2020, Valente et al., 2022]. Critically, because these models are optimized under biological constraints, their learned parameters and dynamics can be interrogated through targeted perturbations to generate testable hypotheses about mechanism. Our study extends this general approach to a well-characterized sensory circuit, applying it to understand the mechanistic basis of a specific feature of neural coding – odor-evoked OFF responses in the insect antennal lobe.

OFF responses, transient increases in firing rate following stimulus termination, are a widespread feature of sensory processing. In the auditory cortex, distinct populations of neurons respond selectively to sound onset and offset [Scholl et al., 2010], and population-level OFF responses have been shown to emerge even when individual neurons exhibit only weak offset activity, playing a functional role in sound termination detection [Bondanelli et al., 2021, Solyga and Barkat, 2021]. In olfaction, OFF responses have been observed across multiple circuit elements: olfactory receptor neurons generate offset response motifs that provide direct feedforward OFF input to downstream circuits [Kim et al., 2023], projection neurons in the locust AL exhibit transient post-stimulus firing increases [Saha et al., 2017], and inhibitory local neurons in the *Drosophila* AL show prominent offset activity that varies with odor identity and concentration [Nagel and Wilson, 2016]. Our RNN model of the AL, which receives realistic ORN inputs including OFF responses at odor offset, reproduces this diversity: PNs exhibit ON, OFF and ON-OFF responses consistent with *in vivo* observations (Fig 2 and [Saha et al., 2017], and LNs in the model also develop offset activity despite not being directly constrained to do so (Fig S7), consistent with the LN offset dynamics reported in *Drosophila* [Nagel and Wilson, 2016].

Despite their prevalence, the mechanisms generating OFF responses in recurrent excitatory-inhibitory circuits remain debated. In the auditory cortex, ON and OFF responses are driven by non-overlapping sets of excitatory synaptic inputs, suggesting that offset activity arises from dedicated feedforward pathways rather than rebound from onset-driven inhibition [Scholl et al., 2010]. By contrast, network modeling of auditory cortex has shown that recurrent dynamics can amplify weak offset signals into robust population-level OFF responses, even without dedicated offset inputs [Bondanelli et al., 2021]. In the olfactory system, in vivo intracellular recordings from the locust AL suggest that disengagement of recurrent inhibition contributes to post-stimulus excitation [Saha et al., 2017]. Computational modeling of the Drosophila AL has similarly shown that the interplay between excitatory and inhibitory connectivity can give rise to diverse temporal motifs, including offset activity in both PNs and LNs [Lazar et al., 2023]. Our model provides direct support for this hypothesized disinhibitory mechanism while revealing that it operates alongside a distinct feedforward pathway.

Decomposition of excitatory and inhibitory inputs onto OFF-responding PNs reveals two mechanistically distinct strategies for generating post-stimulus excitation (Fig 5): in the presence of ORN offset input, both excitatory and inhibitory drive rise at stimulus termination with excitation slightly exceeding inhibition, consistent with a feedforward mechanism driven by dedicated offset inputs. In the absence of offset input, excitation returns to baseline while inhibition drops *below* pre-stimulus levels through LN-LN disinhibition, yielding net excitation through a fundamentally different recurrent mechanism - similar to the release of inhibition pathway proposed by [Saha et al., 2017]. Our model further identifies the specific circuit motif responsible: mutual inhibition among LNs causes a subset to be suppressed slightly below baseline at stimulus termination, releasing their downstream PN targets from tonic inhibition. This provides a concrete mechanistic account of how disengagement of recurrent inhibition generates OFF responses. The two pathways recruit largely non-overlapping neuronal populations, indicating that OFF neuron identity is an emergent property of the network state rather than a cell-intrinsic feature.

A known limitation of data-constrained RNN approaches is that models lacking structural constraints may not faithfully recover the dynamics of unconstrained (unrecorded) neurons or the connectivity of the underlying circuit. Recent theoretical work has shown that connectome-constrained networks can recover unconstrained neuron activity when a sufficient number of neurons are constrained, but that models without synaptic weight constraints may fail to recapitulate unconstrained activity beyond baseline levels, raising concerns about whether such models capture real dynamical features of the original system [Beiran and Litwin-Kumar, 2025]. Similarly, connectivity estimates inferred from neural activity data can be systematically incorrect, particularly in strongly connected recurrent circuits [Das and Fiete, 2020]. Despite these limitations, data-constrained RNNs have been widely and successfully used to infer mechanistic principles across a range of neural systems [Durstewitz et al., 2022, Koppe et al., 2019, Perich et al., 2020, Valente et al., 2022], and we take several steps to mitigate these concerns in our model. We constrain the network architecture to match the known cell-type composition and connectivity of the locust AL (830 PNs and 300 LNs with realistic connection probabilities, Dale’s principle enforcement, and no PN-PN connections) and reject solutions in which unconstrained neuron activity or learned parameters fall outside biologically plausible ranges. The trained model reconstructs constrained PN firing rates with high fidelity (Fig 1), and unconstrained PNs exhibit biologically plausible temporal response patterns and response-type diversity consistent with *in vivo* observations (Fig 2) and [Nagel and Wilson, 2016, Saha et al., 2017]. Importantly, we do not claim to reconstruct the exact time course of every unrecorded neuron or to recover precise synaptic weights. Rather, we focus on the coarse temporal dynamics of the network and the circuit-level mechanisms responsible for generating them. Consistent with this goal, unconstrained LNs in the model develop OFF responses that recapitulate offset dynamics previously recorded in *Drosophila* AL inhibitory interneurons [Nagel and Wilson, 2016], providing additional evidence that the model captures biologically relevant features of the inhibitory population. Finally, the model remains stable and produces interpretable results across all perturbation conditions examined (Figs 3–5), suggesting that the learned dynamics are robust rather than fragile artifacts of overfitting to the constrained neurons.

In summary, our work demonstrates that a biologically-constrained RNN trained directly on *in vivo* recordings can serve as an interpretable model of a sensory circuit, enabling mechanistic dissection of neural phenomena that would be difficult to isolate experimentally. By combining input and connectivity perturbations with excitatory-inhibitory decomposition, we identified two distinct pathways for generating OFF responses in the locust AL - a feedforward mechanism driven by ORN offset input and a recurrent mechanism mediated by LN-LN disinhibition - each recruiting non-overlapping neuronal populations. These findings generate specific, testable predictions: selective silencing of offset-type ORNs should abolish the feedforward pathway while sparing recurrently generated OFF responses, and pharmacological disruption of LN-LN inhibition should selectively eliminate the latter. More broadly, our approach illustrates how data-constrained RNNs can complement traditional hand-tuned models by discovering circuit-level solutions directly from empirical data, and we anticipate that this framework will prove increasingly valuable as large-scale recording techniques continue to expand the scale and resolution of available neural datasets across sensory systems.

## Materials and methods

### *In vivo* experimental procedures

#### Locust husbandry

Locusts (*Schistocerca americana*) were bred, hatched, and raised in lab-maintained colonies. Crowded colonies (200+ locusts) were housed in an incubator set on an automatic 12L:12D light cycle, with daytime temperatures at 36.5°C and night at 25.0°C. Locust diet consisted of fresh-cut grass and wheat germ. Grass was grown in multiple hydroponic plant setups. Locust egg sacks were transferred to a separate incubator (37.7°C) within two days of laying, where they were kept until the first hatchling appeared, then transferred back to the colony incubator.

#### Locust surgery

Post-fifth instar locusts of either sex were selected for experimentation as long as they appeared healthy and had both antennae. For locust surgery, limbs and wings were removed, before the incision sites were sealed with Vet Bond (3M). The locusts were then strapped onto a custom-made platform by securing their abdomens with electrical tape. Locusts were positioned on the platform with their head propped at an angle to ensure easier access to the brain. The antennae were threaded through small pieces of clear tubing with the most distal end protruding out and secured in place atop Batik wax pillars formed beside the head. A two-part, low-VOC epoxy (Devcon No. 14250) was inserted inside the tubing and around the medial and proximal portions of the antennae to prevent them from moving further.

A batik wax bowl was built around the head, starting just above the mandibles and encircling the eyes, and connected to the wax pillars. To keep the brain hydrated as the exoskeleton is removed, the bowl was filled with physiologically balanced locust saline solution at room temperature. The saline solution follows the recipe in [Gilles Laurent et al., 1994], in mM as follows: 140 NaCl, 5 KCl, 5 CaCl_2_, 4 NaHCO_3_, 1 MgCl_2_, 6.3 HEPES, pH 7.0; all chemicals from Sigma-Aldrich Gilles Laurent et al. [1994], Laurent and Naraghi [1994].

A small incision between the antennae was used as the starting point for removing the exoskeleton. Once the exoskeleton was removed between the antennae, above the mandibles, and below the eyes, the head was fully open. The glandular tissue surrounding the brain was removed to expose the ALs of the brain. The gut, which the brain rests on, was severed close to the mandible opening and removed through the abdomen to allow supplemental stabilization of the brain. A small wire platform coated in batik wax was inserted under the brain and positioned to properly adjust and stabilize the brain for electrode insertion before being waxed in place at the intersection of the wax bowl. Finally, a small amount of protease was placed on both ALs to break down the protective sheath surrounding the brain, which was then manually removed with very fine forceps.

#### Odor vial preparation

A panel of five odorants was used in the *in vivo* experiments: hexanal, cis-3-hexenyl acetate, pentanal, propylbenzene, and pure mineral oil as a control. Odors were diluted in 10 mL of mineral oil at 1% vol/vol concentration and covered with a cap. Odor vials were prepared the morning of experimentation and sat for at least two hours before use to ensure plenty of odorant particles accumulated in the headspace.

#### Electrophysiology

Following surgery, locusts were placed within a Faraday cage to reduce noise within the *in vivo* extracellular recordings. A saline drip (10 mL/hr) was connected to the batik wax bowl to replenish lost locust saline solution and to continuously perfuse the brain. A silver-chloride reference wire was placed within the saline bath to complete the circuit. A commercially available 16-channel Neuronexus silicon electrode (A2×2-tet-3mm-150-150-121) was used to record voltage signals from PNs within the AL. Electrodes were electroplated with a gold solution (TivaGlo-Free 24K) for impedance between 200 and 300 kΩ on the morning of experiments.

Using a micromanipulator, electrodes were inserted approximately 100 *µ*m into the AL. Voltage activity signals were sampled at 20 kHz from 8 channels (2 tetrodes). Signals were digitized by an Intan pre-amplifier board (C3334 RHD 16-channel headstage) and transmitted to an Intan recording controller (C3100 RHD USB interface board). Voltage data was recorded through the Intan graphical user interface, where it could also be visualized. LabView data acquisition system automated and synchronized recording time (Intan) and odor delivery (Olfactometer).

Voltage recordings were monitored visually throughout the experiments to ensure no noise artifacts occurred and that the recording remained stable throughout. Each recording lasted approximately 25-30 minutes, and multiple recordings could be done from one locust. It is important to note that data from positions were only recorded when an odor-evoked response was visually seen; if no response was seen, the electrode was backed out and inserted elsewhere until a good position was found. A total of 16 electrophysiological recordings were conducted from 8 locusts.

#### Odor stimulation

This odor stimulation protocol follows previously established methods [Farnum et al., 2023, Liu et al., 2025, McLane-Svoboda et al., 2025, Parnas et al., 2024, Sanchez et al., 2025]. A commercial olfactometer (Aurora Scientific 220A) was used for controlled odorant delivery. Odor vials (five) were attached to each port of the olfactometer using clear 1/16-inch diameter PTFE tubing, with the inlet line going in one side and the outlet line in the other. An odor delivery line connected to the final valve of the olfactometer was set up so that the end of the line was 2-3 cm away from the distal end of one antenna.

At the beginning of each trial, 200 standard cubic centimeters per minute (sccm) of clean air passed through the final valve and onto the antenna with no odor stimulus present. Five seconds before odor delivery, 40% (80 sccm) of air flowing through the olfactometer was redirected into the selected odor vial inlet, picking up the odor headspace, exiting the vial, and rejoining the other 60% (120 sccm) of non-redirected clean air. This was done to prime the lines with the selected odorant before releasing it onto the locust antenna.

When odor stimulation was triggered, the final valve switched from only clean air to the odor + clean air combination and released the stimulus on the antenna. After four seconds of constant odor stimulation, the odor + clean air combination was redirected to the exhaust, and pure clean air flowed through the final valve onto the antenna again. A 6-inch funnel pulling a slight vacuum was placed behind the locust to remove leftover odorants after stimulation. This protocol maintains constant air flow onto the antenna to eliminate any neural responses due to mechanosensory detection of air pressure changes. For each recording position, the order of odor stimulus was pseudo-randomized to prevent accumulating bias of recording signal over time. For each experiment, all five odors were presented to the locust antenna for four seconds, five trials, with a one-minute inter-stimulus interval.

#### Projection neuron spike sorting

Neural data was passed through a high-pass 300 Hz Butterworth filter to eliminate low frequencies and then imported into MATLAB. Using a custom-written MATLAB R2022a code, the data was transformed into a readable Igor Pro 4.06A format. Spike sorting analysis was performed in Igor following previously described methods [Pouzat et al., 2002]. Briefly, detection thresholds were set between 2.5 and 3.5 standard deviations (SD) of baseline to find neural spikes.

The initial criteria for individual PNs to pass were cluster separation (nearest projection) ≥ 5 SD, inter-spike intervals (ISIs) ≤ 10%, and spike waveform variance (SD test) ≤ 10%. Raster plots of all odors and trials were created from neurons that passed the initial criteria. These raster plots were inspected visually to determine if they met the secondary criteria: consistent spiking baseline across trials and odors, and no portions of trials were missing. A total of 110 spike sorted neurons from 9 locusts passed both criteria. Spike sorted data was used for all analysis and RNN training.

#### Dimensionality reduction analysis

Three separate data groups were analyzed using principal component analysis (PCA) to visualize odor-evoked neural activity across the population of neurons. The population of neurons collected was pooled over all electrophysiology experiments for analysis. The first group of data consisted of 110 PNs collected from the locust AL *in vivo*; the second group was recurrent neural network (RNN) PNs that were trained to replicate the firing rates of the 110 *in vivo* PNs; the third group was all 830 RNN PNs.

Firing rates created from spike sorted data in each group were smoothed using a moving window average of 25 ms bins, incrementing by 5 ms. Data was then transformed into 50 ms non-overlapping bins, aligning with the 20 Hz oscillations found in the locust AL, and the activity was normalized by subtracting the average firing rate of one second before stimulus onset. The 4-D matrix unique to each data group consisted of dimensions: neurons × time bins × trials × odors, where each element holds the firing rate of one neuron in one 50 ms time bin. All five trials of one odor were then averaged. For example, spike sorted and binned neural responses for a specific odor in the *in vivo* group generated a matrix of neuron number (110) × time bins (140, number of 50 ms bins between -1 to 6 s). In the end, three 4-D matrices corresponding to each of the groups were created and used for the following analyses.

In PCA, the high-dimensional vector in each time bin was projected along the eigenvectors of the covariance matrix, plotted within the high-dimensional space. This was then reduced to three dimensions by taking the highest eigenvalues that maximize variance within the dataset and plotted. Adjacent time bins were connected to visualize the population neural response for each odor over time. Neural trajectories were smoothed using a third-order IIR Butterworth filter (Half Power Frequency = 0.15). Each odor trajectory was shifted to begin at the origin for easier visualization of response dynamics and separation. All analysis was performed using custom-written code in MATLAB R2022a.

### Antennal lobe circuit model

We constructed a firing rate-based recurrent neural network (RNN) model of the locust antennal lobe (AL) comprising two neuron populations: 830 excitatory projection neurons (PNs) and 300 inhibitory local interneurons (LNs), receiving input from 1,000 olfactory receptor neurons (ORNs). The network architecture was constrained to reflect known AL connectivity. ORN inputs projected to both PNs and LNs with full connectivity (connection probability *p* = 1.0). Among recurrent connections, the PN → LN, LN → PN, and LN → LN pathways were each connected with a probability of *p* = 0.4; no PN → PN connections were present (*p* = 0.0). Self-connections were excluded. Binary connectivity masks were generated by drawing Bernoulli random variables at the specified connection probabilities and remained fixed throughout training.

#### ORN input

Olfactory receptor neurons (ORNs) provided excitatory input to both PN and LN populations. All ORNs were connected to all PNs and LNs, though with heterogeneous synaptic strengths. Synaptic weights were scaled to produce physiologically realistic postsynaptic responses (ORN → PN: stronger drive; ORN → LN: moderate drive).

The ORN activity was generated using a MAP based model and recreated the different temporal motifs observed in locust ORN activity [Kim et al., 2023] (Fig S2A). The different odor inputs to the MAP based model were designed to capture the chemical similarity between Hexanal and the other odors (Fig S2B, C). The MAP model generates spiking activity for each odor and trial, which was then converted to firing rates and used as input to the RNN model.

#### Neuron dynamics

Each neuron’s state variable *x*_*i*_ evolved according to a first-order ordinary differential equation, discretized with an Euler integration step of Δ*t* = 5 ms:

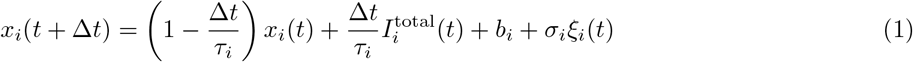

where *τ*_*i*_ is the neuron’s time constant, *I*^total^ is the total synaptic input current, *b*_*i*_ is a bias term, *σ*_*i*_ is the noise scale, and *ξ*_*i*_(*t*) ∼ 𝒩 (0, 1) is independent Gaussian noise at each time step. The instantaneous firing rate was obtained by passing the state variable through a clipped softplus activation function:

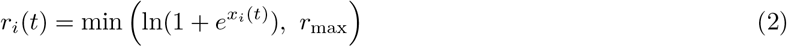

where *r*_max_ = 120 Hz, reflecting physiological constraints on maximum firing rates observed in locust AL neurons.

#### Time constants

Neuron time constants were heterogeneous across the population and trainable via backpropagation. They were parameterized using a sigmoid transformation to enforce biologically plausible bounds:

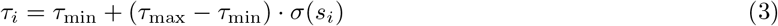

where *s*_*i*_ is a learnable parameter and *σ*(·) is the logistic sigmoid function. The bounds were set to *τ*_min_ = 15 ms and *τ*_max_ = 300 ms for both PNs and LNs. The learnable parameters *s*_*i*_ were initialized from a log-normal distribution for both neuron types.

#### Bias and noise

Each neuron received a trainable bias current drawn independently from a Gaussian distribution at initialization: 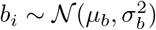. For PNs, *µ*_*b*_ = 0.0 and *σ*_*b*_ = 0.01; for LNs, *µ*_*b*_ = 0.0 and *σ*_*b*_ = 0.02. Additive Gaussian noise was applied at each time step with scale *σ* = 0.01 for PNs and *σ* = 0.02 for LNs.

#### Synaptic connectivity and Dale’s principle

All recurrent synaptic connections obeyed Dale’s principle: PN outputs were constrained to be excitatory (non-negative effective weights) and LN outputs were constrained to be inhibitory (non-positive effective weights). This constraint was enforced by taking the absolute value of the raw weight parameters and applying the appropriate sign:

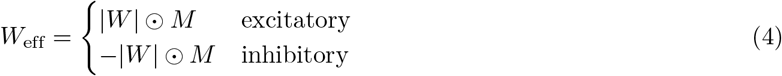

where *M* is the binary connectivity mask and ⊙ denotes element-wise multiplication. Input synapses (ORN→PN and ORN→LN) were excitatory and initialized as *W*_*ij*_ = |*w*_*ij*_| where *w*_*ij*_ ∼ 𝒩 (0, *α*^2^*/N*_pre_) with *N*_pre_ = 1000; the scaling factor was *α* = 2.5 for ORN→PN and *α* = 1.0 for ORN→LN. Recurrent synapses were initialized as 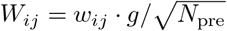 where *w*_*ij*_ ∼ 𝒩 (0, 1), then masked by the binary connectivity matrix and constrained by Dale’s principle. The weight scale was *g* = 2.0 for the PN→LN, LN→PN, and LN→LN pathways.

#### Synaptic current computation

At each time step, the total input current to each neuron was computed as the sum of feedforward and recurrent contributions:

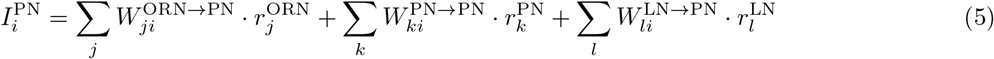

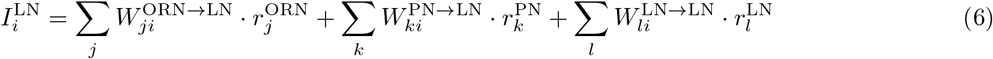

where the PN → PN recurrent current was zero due to the absence of those connections (*p* = 0). The remaining recurrent pathways (PN → LN, LN→ PN, and LN → LN) all contributed non-zero currents with trainable weights. All neuron state variables and firing rates were initialized to zero at the start of each trial (*x*_*i*_(0) = 0, *r*_*i*_(0) = 0), and the network was driven into activity solely by the ORN input.

#### Training data

The model was trained on experimentally recorded data from the locust AL. Firing rates were smoothed with a Gaussian kernel. Each trial consisted of a 7-second simulation (1,400 time bins at 5 ms resolution) with the following temporal structure: a 1-second pre-stimulus baseline (bins 0–200), a 4-second odor stimulus (bins 200–1000), and a 2-second post-stimulus period (bins 1000–1400).

#### Loss function and optimization

The total training loss combined a reconstruction term with regularization terms that constrained the activity of unrecorded neurons:

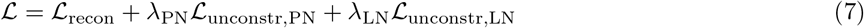

The reconstruction loss was the root mean squared error (RMSE) between the model’s PN firing rates and the experimentally recorded PN firing rates over all time steps:

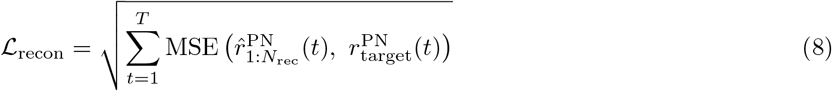

where *N*_rec_ = 110 is the number of experimentally recorded PNs (out of 830 total model PNs) and *T* is the number of time steps. The remaining 720 unrecorded PNs were not directly constrained by the reconstruction loss. To prevent unconstrained neurons from developing unrealistic activity, the mean firing rate of a randomly selected subset (75%) of unconstrained PNs and LNs was penalized to match the mean of the recorded PN target activity, computed over non-overlapping windows of 200 time steps:

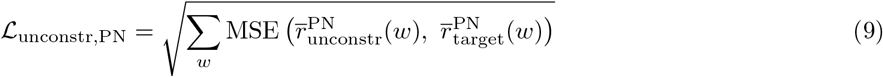

with an analogous term for LNs. The regularization weights were *λ*_PN_ = *λ*_LN_ = 0.15.

The model was trained using the Adam optimizer with AMSGrad for 6,000 epochs, with an initial learning rate of 7.5 × 10^−4^ and an exponential decay schedule (*γ* = 0.9995 per epoch). Mixed-precision training (float16 autocast with gradient scaling) was employed for computational efficiency, with a batch size of 10 sequences.

Trainable parameters included all recurrent synaptic weights (PN → LN, LN → PN, and LN → LN), input synaptic weights (ORN → PN and ORN → LN), and all neuron time constants for both PNs and LNs. Input synaptic weights were made trainable from the middle of training and remained trainable for a fixed number of epochs.

### Analysis of RNN model responses

#### Input perturbation

To assess the contribution of specific ORN subpopulations to PN response dynamics, we selectively perturbed ORN inputs to the trained model. The firing rates of offset-type ORNs were multiplicatively scaled by a perturbation factor *α*:

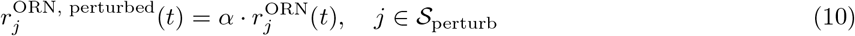

where 𝒮_perturb_ is the set of perturbed ORN indices (i.e., ORNs exhibiting offset-type responses) and *α* ∈ {0.0, 0.25, 0.5, 0.75, 1.0, 1.25, 1.5, 1.75, 2.0}. A factor of *α* = 1.0 corresponds to the intact (baseline) network, *α* = 0.0 removes the selected ORN inputs entirely, and *α >* 1.0 amplifies them. ORNs outside 𝒮 _perturb_ were left unchanged, ensuring that only the offset component of the input was manipulated. All other network parameters (recurrent weights, time constants, biases) remained fixed at their trained values.

For each perturbation factor, the trained model was simulated using the perturbed ORN input, and PN and LN firing rate trajectories were recorded over the full 7-second trial duration across all odors and trials. For each perturbation condition, PN response type counts were tallied using the classification procedure described above, E/I input decomposition was computed to assess shifts in excitatory-inhibitory balance across PN response-type groups, and population-averaged PN and LN firing rates were computed across all odors, trials, time bins, and neurons. This approach allowed us to isolate the contribution of direct ORN offset input to the generation and functional role of PN OFF responses.

#### Network perturbation

To dissect the functional role of inhibitory circuitry in the AL network, we systematically scaled the trained inhibitory synaptic weights by a factor *β* and re-simulated the network. Four perturbation types were examined: (1) Global inhibition, scaling both LN → LN and LN → PN weights; (2) LN-LN inhibition, scaling LN → LN weights only; (3) LN-PN inhibition, scaling LN → PN weights only; and (4) PN-LN excitation, scaling PN → LN weights only. For each perturbation type, the raw weight parameters of the specified synaptic pathway(s) were scaled:

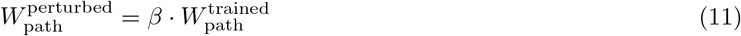

with *β* ∈ { 0.0, 0.25, 0.5, 0.75, 1.0, 1.25, 1.5, 1.75, 2.0} . A factor of *β* = 1.0 corresponds to the intact network, *β* = 0.0 ablates the pathway entirely, and *β >* 1.0 strengthens it. Dale’s principle constraints remained enforced through the effective weight computation (i.e., the sign-constrained absolute-value parameterization was preserved). All other network parameters were held fixed.

For each perturbation type and factor, the number of ON, OFF, and ON-OFF PNs was tallied using the classification criteria described above and compared to the intact network baseline. Additionally, excitatory and inhibitory input currents to each PN were computed via the E/I decomposition and grouped by PN response type to assess how each pathway perturbation alters the excitatory-inhibitory balance during OFF time period.

#### Classifying neuron responses

Neurons (*in vivo* and model PNs both) were sorted into one of four different responses for each odor trial: onset, offset, onset-offset, and no response. Baseline activity was defined as the second before odor stimulus (-1 to 0 sec), therefore activity within that timeframe had no odor specific information. Onset and offset activity windows, respectively, were defined as the four seconds during odor presentation (0 to 4 sec) and the two seconds after (4 to 6 sec). For each neuron, the average firing rate and standard deviation (SD) were calculated from baseline activity pooled from every trial regardless of odor. Neuron specific activity thresholds (AT) were calculated by taking the average baseline firing rate + 2SD. Additionally, a minimum threshold (MT) of 5 Hz was included in our criteria to eliminate neurons that had very little activity but still passed the AT due to low baseline values.

Using the parameters above, responses were classified based on individual neural activity for every odor and trial. Neurons for a specific odor trial were marked as an onset response only if the average activity during odor onset window exceeded the AT, but the activity during the offset window did not, and the peak firing rate during onset was above the MT. Average activity during each activity window was calculated using a moving 500 msec window. Responses were sorted as offset only if the average activity in the odor offset window exceeded the threshold, but the onset activity did not, and the peak firing rate during offset passed the MT. Responses were sorted as onset-offset if the activity in both time windows passed the AT and MT thresholds. Lastly, responses were classified as having no response if the activity during onset and offset did not pass either AT or MT criteria. This sorting was repeated for the same neuron across all odors and trials before continuing to every neuron after. Results were displayed showing trial variability and separated based on odor to visualize trends in odor evoked responses.

#### Excitatory/inhibitory balance analysis

To characterize the balance of excitatory and inhibitory drive onto model PNs, we decomposed the total synaptic input into excitatory and inhibitory components at each time step. For each PN *i*, the excitatory input was computed as the product of the trained ORN→PN weight matrix and the ORN firing rate vector, and the inhibitory input as the product of the absolute LN→PN weight matrix and the LN firing rate vector:

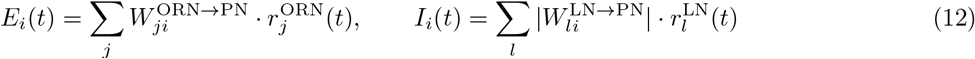

These quantities were then averaged within PN response-type groups (ON, OFF, ON-OFF, non-responsive) to assess how perturbations alter the excitatory-inhibitory balance across functionally distinct PN subpopulations. This decomposition was applied to both input perturbation and inhibitory pathway perturbation analyses.

## Acknowledgments

MGNZ received support form the Schmidt Sciences Foundation. AKMS received funding support from National Science Foundation Graduate Research Fellowship. MB received funding support from National Science Foundation (BCS-2323241 and EFMA-2223839) and National Institutes of Health (1R01DC020892). DS received funding from National Science Foundation (2323240).

## Supporting information

**Fig S1.**
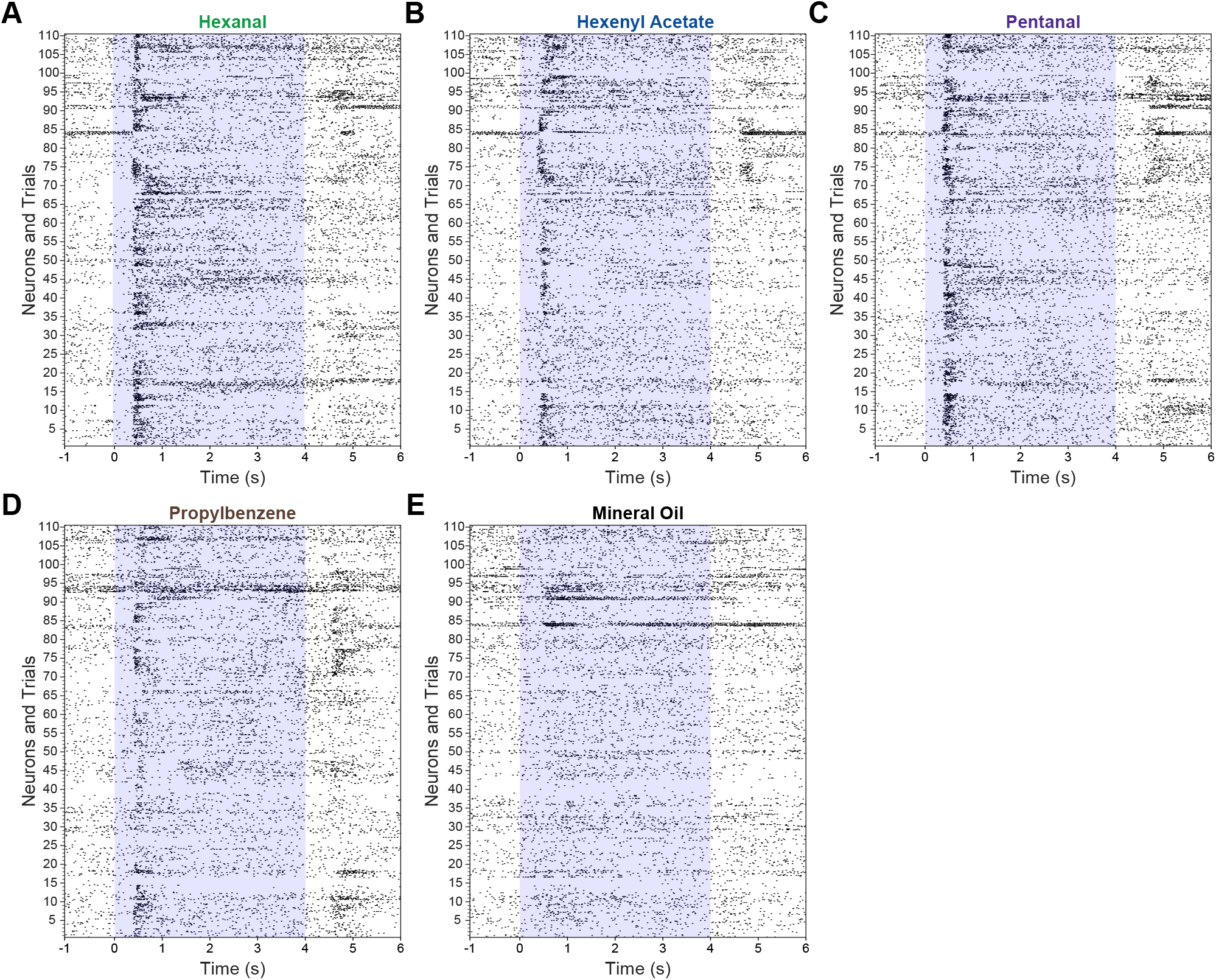
*In vivo* PN raster plots. A) Spiking events for hexanal. Each black line represents an action potential; every row is one trial from one neuron. All 110 *in vivo* neurons are shown with 5 trials each. Shaded box is odor stimulus of 4 seconds. Raster plot shows an increase in firing during stimulus onset over the majority of neurons. B) Odor-evoked response for each neuron to hexenyl acetate shows an increase in spiking during odor presentation. C) Activity to pentanal shows an increase during odor stimulus and more neurons firing after odor is turned off. D) Activity over neurons shows a weaker increase in firing in response to propylbenzene onset, but more neurons having an offset response compared to other odors. E) Mineral oil has little affect on individual neuron spiking.

**Fig S2.**
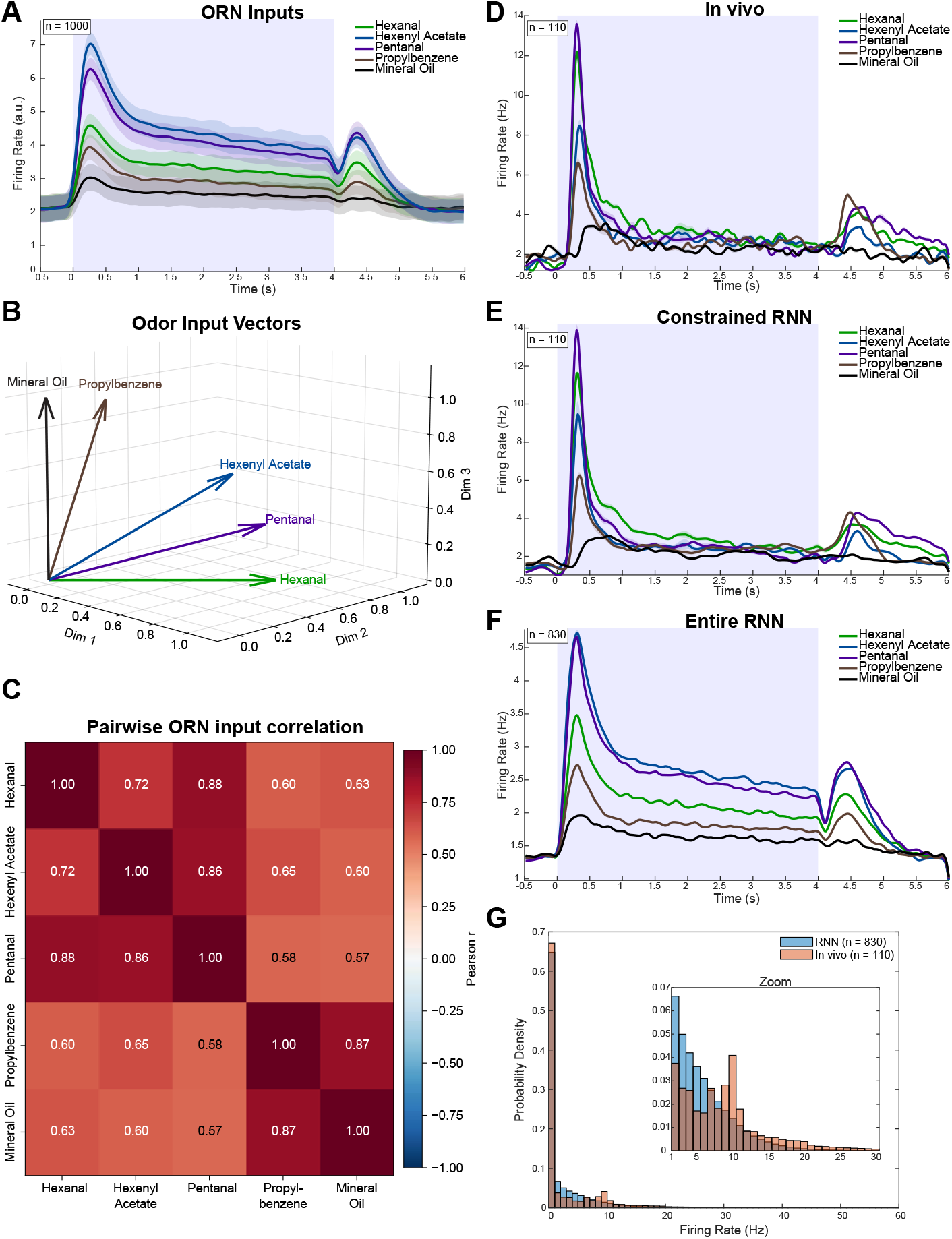
ORN inputs and comparison of in vivo and model PN populations. A) ORN input firing rates (n = 1000 simulated ORNs) for each of the five odorants, averaged across the population. Shaded regions around average firing rate indicate SEM and the blue box marks the odor stimulus window (0 to 4 seconds). B) Odor input vectors in three-dimensional input space. Each vector represents the input pattern used to drive ORN activity for a single odorant, and the angle between vectors reflects the similarity of the corresponding odor representations, with smaller angles indicating more similar inputs. These inputs were designed to capture the chemical similarity between hexanal and the remaining odorants. C) Pairwise correlation between ORN input patterns across odors, quantifying the similarity of ORN-level odor representations. Chemically related odors (e.g., hexanal and pentanal) show higher correlations. D) Population peri-stimulus time histogram (PSTH) of odor-evoked firing rates for all in vivo PNs (n = 110). Odor is delivered to the antennae at 0 seconds and turned off at 4 seconds. Each odorant evokes a distinct temporal response profile, with increased firing at stimulus onset and offset. E) PSTH for constrained model PNs (n = 110), which closely match the odor-evoked responses of their in vivo counterparts in D. F) PSTH for all model PNs (constrained and unconstrained, n = 830), showing that population-level temporal dynamics generalize across the full PN population. G) Probability density showing firing rate distribution for in vivo responses (peach) compared to the entire RNN (blue). Though slightly different, RNN firing rates are within biologically plausible ranges, matching in vivo data. Inset shows a zoomed view of the lower firing rate range.

**Fig S3.**
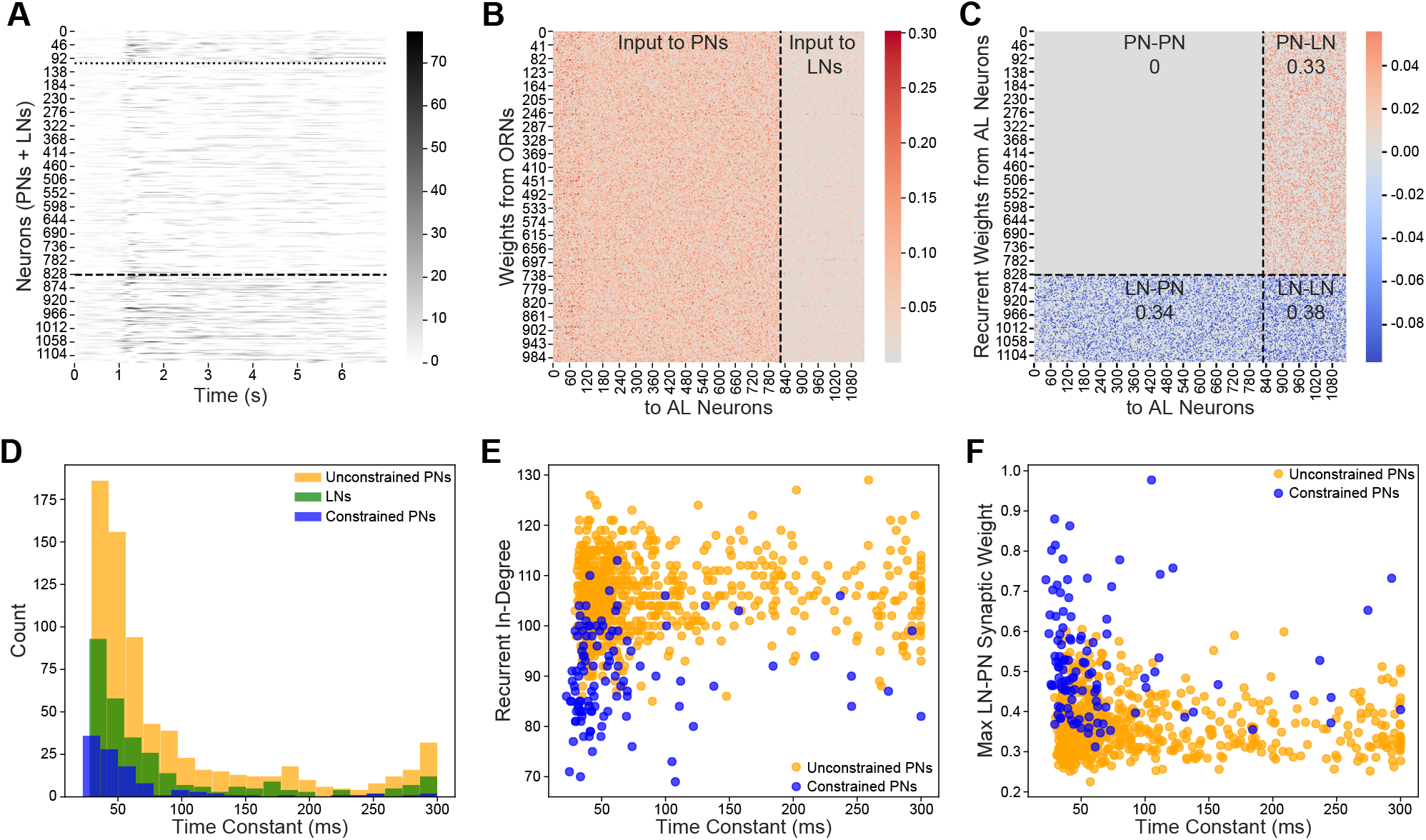
Parameters of the optimized AL model. A) Firing rates of all neurons (PNs + LNs in Hz) in response to hexanal stimulation, trial 1. Dotted line delineates constrained and unconstrained PNs; dashed line separates PNs from LNs. B) Heatmap of optimized input connectivity from 1000 ORNs (y-axis) to 830 PNs and 300 LNs (x-axis). Input weights were initialized with stronger connectivity to PNs relative to LNs. C) Heatmap of optimized recurrent connectivity within the AL network. Dashed lines delineate PNs and LNs. Excitatory connections are shown in red, inhibitory connections in blue. Connection densities for each quadrant are indicated. Connection densities were initialized to be 0.4 before optimization. D) Distribution of optimized time constants (ms) for constrained PNs (blue), unconstrained PNs (orange), and LNs (green). E) Relationship between recurrent in-degree and optimized time constants for PNs. Constrained PNs exhibit lower in-degrees and shorter time constants compared to unconstrained PNs. F) Relationship between maximum inhibitory synaptic weight and time constant for PNs in the optimized model. Constrained PNs consistently receive stronger inhibitory inputs from LNs than unconstrained PNs.

**Fig S4.**
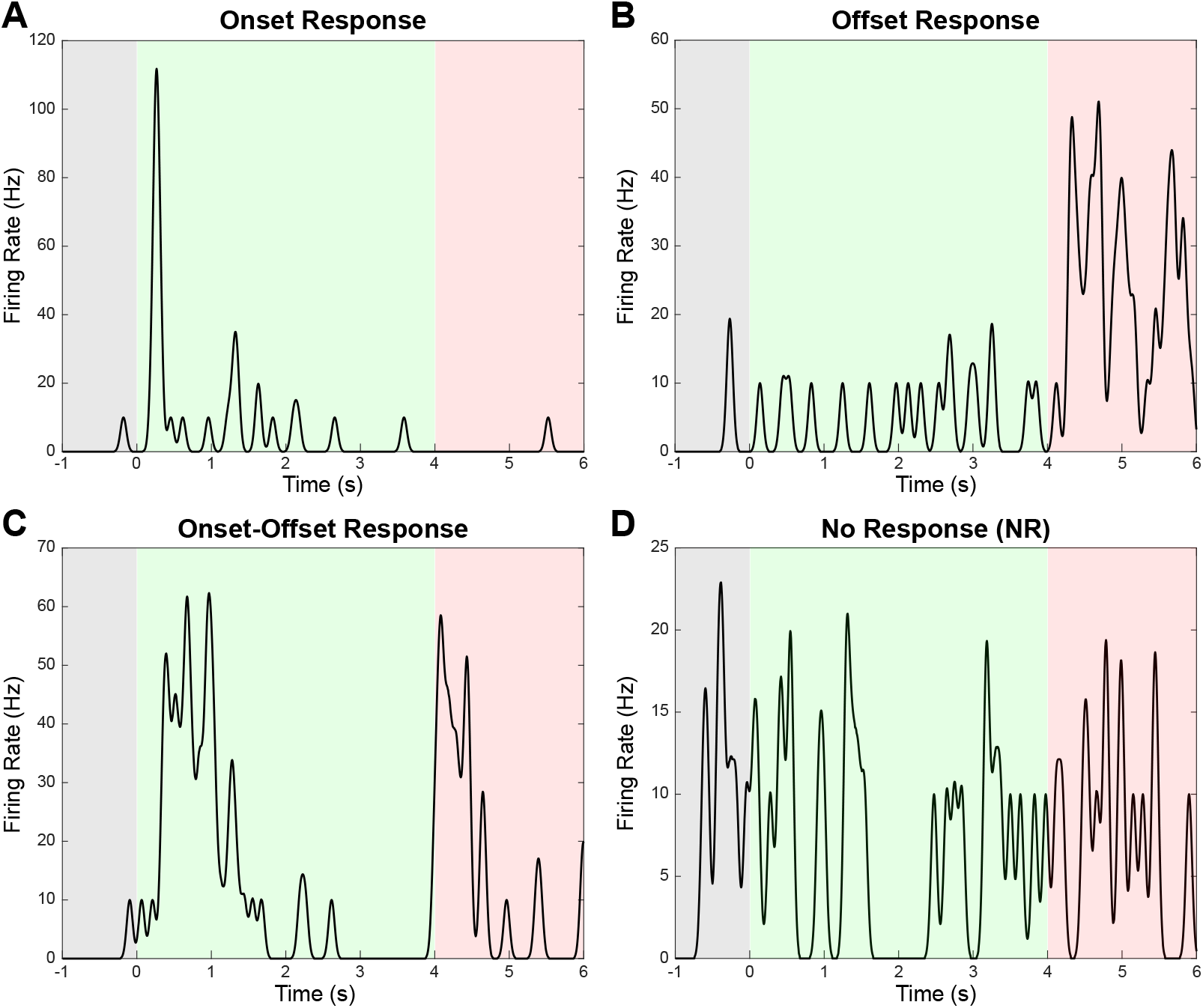
Representative examples for each type of response classification. A) Peristimulus time histogram (PSTH) showing a response classified as onset. Onset activity was defined as activity passing a set criteria within the odor stimulus time frame (green box, see Methods for criteria). Graph depicts the response from one neuron, one odor, one trial. The gray box is baseline activity (before odor stimulus). B) PSTH showing an offset response, defined as activity exceeding set criteria after the odor stimulus is turned off (red box). C) PSTH showing an onset-offset response where neural activity passed the criteria for both onset and offset time frames. D) Representative neural activity that was classified as no response, as it had no change in onset or offset activity compared to baseline.

**Fig S5.**
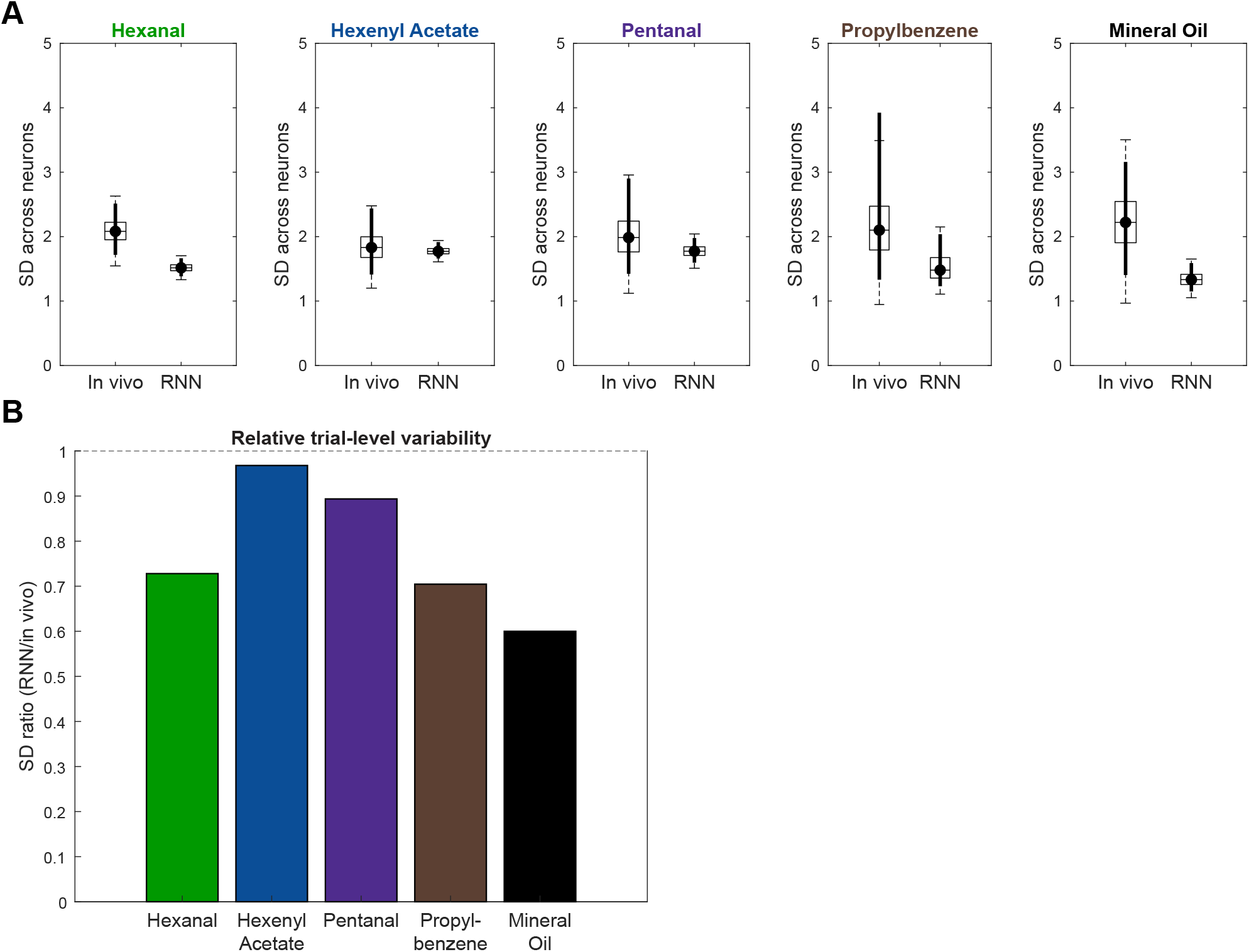
Trial-level variability across neurons. A) Standard deviation (SD) of firing rate across neurons separated by odor. Estimated using bootstrap resampling to account for unequal neuron total (110 *in vivo* vs 830 RNN). Box and whiskers show bootstrap distributions, circles indicate medians, bold vertical lines denote 95% confidence intervals (CI). Medians were similar between *in vivo* and RNN datasets across all odors, with bootstrap CIs overlapping. B) SD ratios were calculated by taking the median CI of the bootstrapped RNN and dividing it by the median CI of the bootstrapped *in vivo* data. Ratios remained between 0.6 and 1 across odors, indicating that the datasets are comparable in terms of neural activity, with ratios significantly greater or less than one being expected if one dataset had higher variability in neural activity between trials.

**Fig S6.**
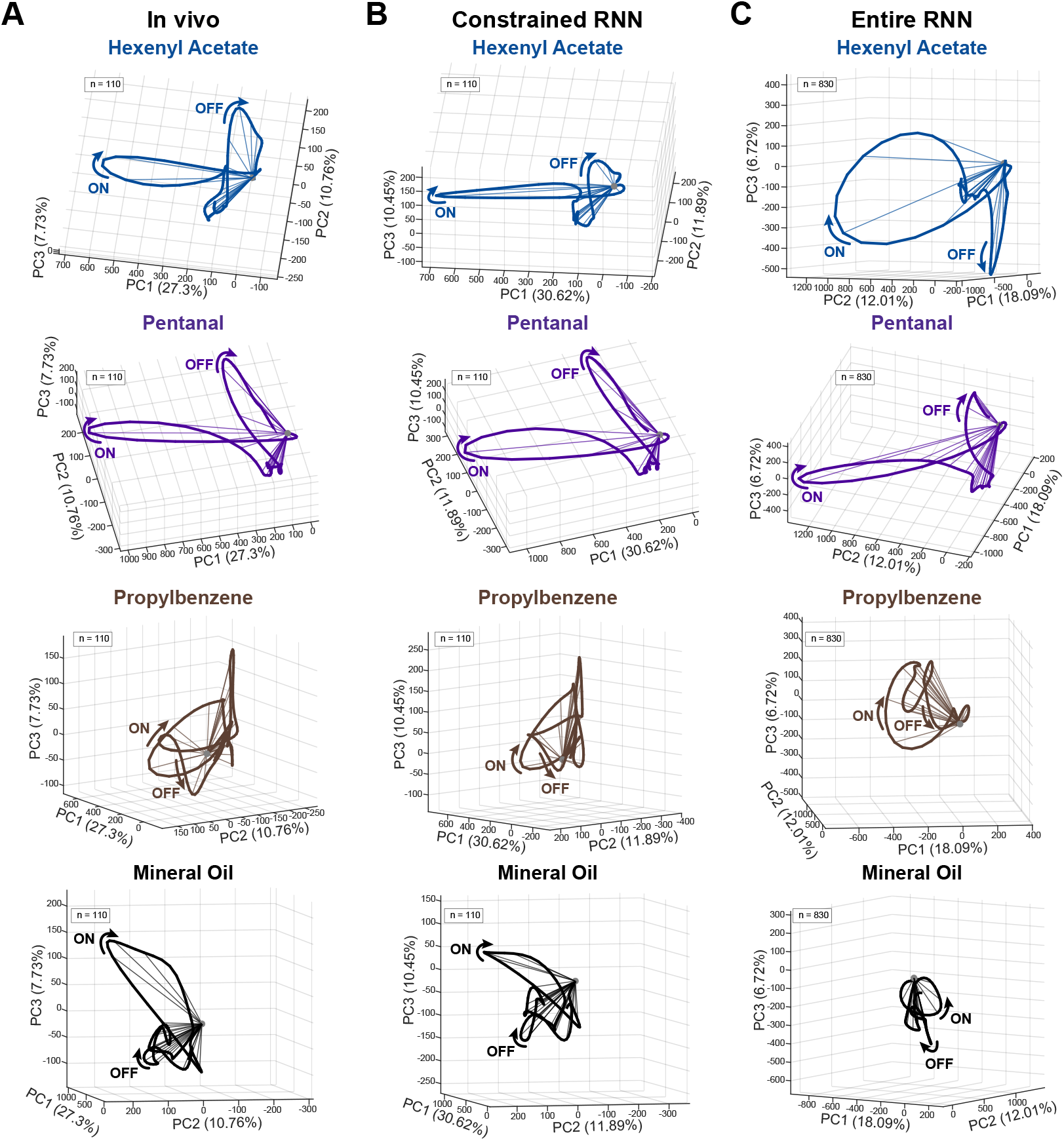
Temporal dynamics for the additional four odors. A) Principal component analysis (PCA) of *in vivo* PNs (*n* = 110) showing neural trajectories over time for the other four odors: hexenyl acetate (blue), pentanal (purple), propylbenzene (brown), and mineral oil (black). Distinct onset (labeled on) and offset (labeled OFF) trajectory loops can be seen. See Fig 2 for hexanal PCAs. B) PCAs for constrained model PNs (*n* = 110), with comparable odor-specific trajectories and looping patterns as seen in *in vivo*. C) PCAs of all model PNs (constrained and unconstrained, *n* = 830). The entire RNN displays temporal properties similar to those of the recorded *in vivo* PNs. -0.5 to 6 seconds is displayed in the PCAs, with odor onset at 0 and offset at 4 seconds.

**Fig S7.**
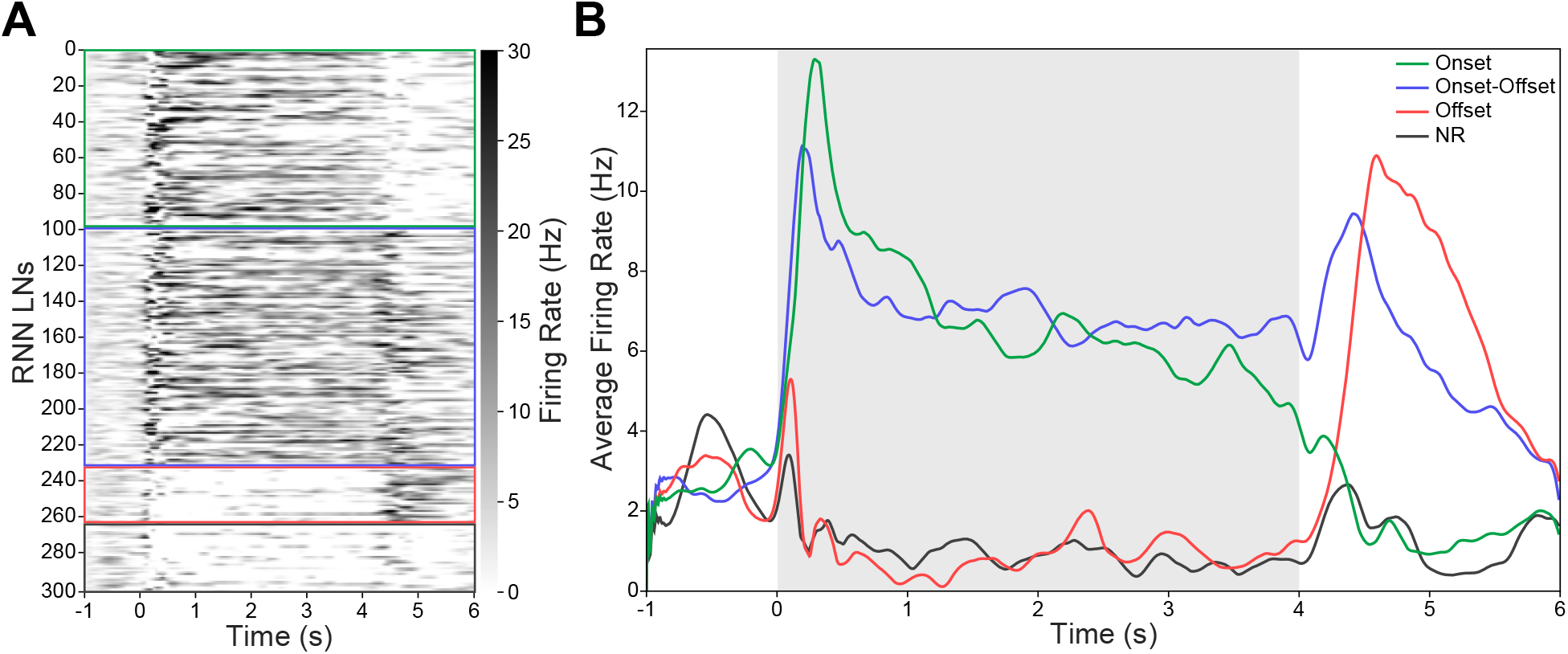
OFF responses in LNs. A) Firing rate heatmap of all 300 RNN LNs for one trial of a representative odor. Odor stimulus is delivered from 0 to 4 seconds. LNs are sorted by response type, with colored lines delineating onset, onset-offset, offset, and non-responsive groups (classified using the same criteria applied to PNs; see Methods). B) Average firing rate profiles of onset (green), onset-offset (blue), offset (red), and non-responsive (NR, black) LNs in response to stimulation (representative odor shown). Gray shaded region indicates the odor stimulus window (0 to 4 seconds). A subpopulation of LNs exhibits OFF responses, with firing rates that rise following stimulus termination.

## References

Omri Barak. Recurrent neural networks as versatile tools of neuroscience research. Current Opinion in Neurobiology, 46:1–6, October 2017. ISSN 09594388. doi: 10.1016/j.conb.2017.06.003. URL https://linkinghub.elsevier.com/retrieve/pii/S0959438817300429.

Maxim Bazhenov, Mark Stopfer, Mikhail Rabinovich, Ramon Huerta, Henry D.I. Abarbanel, Terrence J. Sejnowski, and Gilles Laurent. Model of Transient Oscillatory Synchronization in the Locust Antennal Lobe. Neuron, 30(2):553–567, May 2001. ISSN 08966273. doi: 10.1016/S0896-6273(01)00284-7. URL https://linkinghub.elsevier.com/retrieve/pii/S0896627301002847.

Maxim Bazhenov, Mark Stopfer, Terrence J. Sejnowski, and Gilles Laurent. Fast Odor Learning Improves Reliability of Odor Responses in the Locust Antennal Lobe. Neuron, 46(3):483–492, May 2005. ISSN 0896-6273. doi: 10.1016/j.neuron.2005.03.022. URL https://www.ncbi.nlm.nih.gov/pmc/articles/PMC2905210/.

Maxim Bazhenov, Ramon Huerta, and Brian H. Smith. A Computational Framework for Understanding Decision Making through Integration of Basic Learning Rules. The Journal of Neuroscience, 33(13):5686–5697, March 2013. ISSN 0270-6474. doi: 10.1523/JNEUROSCI.4145-12.2013. URL https://www.ncbi.nlm.nih.gov/pmc/articles/PMC3667960/.

Manuel Beiran and Ashok Litwin-Kumar. Prediction of neural activity in connectome-constrained recurrent networks. Nature Neuroscience, pages 1–14, October 2025. ISSN 1546-1726. doi: 10.1038/s41593-025-02080-4. URL https://www.nature.com/articles/s41593-025-02080-4.

Giulio Bondanelli, Thomas Deneux, Brice Bathellier, and Srdjan Ostojic. Network dynamics underlying OFF responses in the auditory cortex. eLife, 10:e53151, March 2021. ISSN 2050-084X. doi: 10.7554/eLife.53151. URL https://elifesciences.org/articles/53151.

Jen-Yung Chen, Emiliano Marachlian, Collins Assisi, Ramon Huerta, Brian H. Smith, Fernando Locatelli, and Maxim Bazhenov. Learning Modifies Odor Mixture Processing to Improve Detection of Relevant Components. The Journal of Neuroscience, 35(1):179–197, January 2015. ISSN 0270-6474, 1529-2401. doi: 10.1523/JNEUROSCI.2345-14.2015. URL https://www.jneurosci.org/lookup/doi/10.1523/JNEUROSCI.2345-14.2015.

Abhranil Das and Ila R. Fiete. Systematic errors in connectivity inferred from activity in strongly recurrent networks. Nature Neuroscience, 23(10):1286–1296, October 2020. ISSN 1097-6256, 1546-1726. doi: 10.1038/s41593-020-0699-2. URL https://www.nature.com/articles/s41593-020-0699-2.

Laura N. Driscoll, Krishna Shenoy, and David Sussillo. Flexible multitask computation in recurrent networks utilizes shared dynamical motifs. Nature Neuroscience, pages 1–15, July 2024. ISSN 1546-1726. doi: 10.1038/s41593-024-01668-6. URL https://www.nature.com/articles/s41593-024-01668-6.

Alexis Dubreuil, Adrian Valente, Manuel Beiran, Francesca Mastrogiuseppe, and Srdjan Ostojic. The role of population structure in computations through neural dynamics. Nature Neuroscience, 25(6):783–794, June 2022. ISSN 1097-6256, 1546-1726. doi: 10.1038/s41593-022-01088-4. URL https://www.nature.com/articles/s41593-022-01088-4.

Daniel Durstewitz, Georgia Koppe, and Max Ingo Thurm. Reconstructing Computational Dynamics from Neural Measurements with Recurrent Neural Networks, November 2022. URL https://www.biorxiv.org/content/10.1101/2022.10.31.514408v1. Pages: 2022.10.31.514408 Section: New Results.

Alexander Farnum, Michael Parnas, Ehsanul Hoque Apu, Elyssa Cox, Nöel Lefevre, Christopher H. Contag, and Debajit Saha. Harnessing insect olfactory neural circuits for detecting and discriminating human cancers. Biosensors and Bioelectronics, 219:114814, January 2023. ISSN 0956-5663. doi: 10.1016/j.bios.2022.114814. URL https://www.sciencedirect.com/science/article/pii/S0956566322008545.

Gilles Laurent, Gilles Laurent, and Hananel Davidowitz. Encoding of olfactory information with oscillating neural assemblies. Science, 265(5180):1872–1875, September 1994. doi: 10.1126/science.265.5180.1872. MAG ID: 1963860447.

Shruti Joshi, Seth Haney, Zhenyu Wang, Fernando Locatelli, Hong Lei, Yu Cao, Brian Smith, and Maxim Bazhenov. Plasticity in inhibitory networks improves pattern separation in early olfactory processing. Communications Biology, 8(1):590, April 2025. ISSN 2399-3642. doi: 10.1038/s42003-025-07879-2. URL https://www.nature.com/articles/s42003-025-07879-2.

Brian Kim, Seth Haney, Ana P Milan, Shruti Joshi, Zane Aldworth, Nikolai Rulkov, Alexander T Kim, Maxim Bazhenov, and Mark A Stopfer. Olfactory receptor neurons generate multiple response motifs, increasing coding space dimensionality. eLife, 12:e79152, January 2023. ISSN 2050-084X. doi: 10.7554/eLife.79152. URL https://elifesciences.org/articles/79152.

Robert Kim and Terrence J. Sejnowski. Strong inhibitory signaling underlies stable temporal dynamics and working memory in spiking neural networks. Nature Neuroscience, 24(1):129–139, January 2021. ISSN 1546-1726. doi: 10.1038/s41593-020-00753-w. URL https://www.nature.com/articles/s41593-020-00753-w. Number: 1.

Robert Kim, Yinghao Li, and Terrence J. Sejnowski. Simple framework for constructing functional spiking recurrent neural networks. Proceedings of the National Academy of Sciences, 116(45):22811–22820, November 2019. doi: 10.1073/pnas.1905926116. URL https://www.pnas.org/doi/abs/10.1073/pnas.1905926116.

Georgia Koppe, Hazem Toutounji, Peter Kirsch, Stefanie Lis, and Daniel Durstewitz. Identifying nonlinear dynamical systems via generative recurrent neural networks with applications to fMRI. PLOS Computational Biology, 15(8):e1007263, August 2019. ISSN 1553-7358. doi: 10.1371/journal.pcbi.1007263. URL https://dx.plos.org/10.1371/journal.pcbi.1007263.

Janne K. Lappalainen, Fabian D. Tschopp, Sridhama Prakhya, Mason McGill, Aljoscha Nern, Kazunori Shinomiya, Shin-ya Takemura, Eyal Gruntman, Jakob H. Macke, and Srinivas C. Turaga. Connectome-constrained networks predict neural activity across the fly visual system. Nature, 634(8036): 1132–1140, October 2024. ISSN 0028-0836, 1476-4687. doi: 10.1038/s41586-024-07939-3. URL https://www.nature.com/articles/s41586-024-07939-3.

G Laurent and M Naraghi. Odorant-induced oscillations in the mushroom bodies of the locust. The Journal of Neuroscience, 14(5):2993–3004, May 1994. ISSN 0270-6474, 1529-2401. doi: 10.1523/JNEUROSCI.14-05-02993.1994. URL https://www.jneurosci.org/lookup/doi/10.1523/JNEUROSCI.14-05-02993.1994.

Gilles Laurent. Olfactory network dynamics and the coding of multidimensional signals. Nature Reviews Neuroscience, 3(11):884–895, November 2002. ISSN 1471-0048. doi: 10.1038/nrn964. URL https://www.nature.com/articles/nrn964. Number: 11.

Gilles Laurent, Michael Wehr, and Hananel Davidowitz. Temporal representations of odors in an olfactory network. Journal of Neuroscience, 16(12):3837–3847, 1996. ISSN 0270-6474. doi: 10.1523/JNEUROSCI.16-12-03837.1996. URL https://www.jneurosci.org/content/16/12/3837.

Aurel A. Lazar, Tingkai Liu, and Chung-Heng Yeh. The functional logic of odor information processing in the Drosophila antennal lobe. PLOS Computational Biology, 19(4):e1011043, April 2023. ISSN 1553-7358. doi: 10.1371/journal.pcbi.1011043. URL https://dx.plos.org/10.1371/journal.pcbi.1011043.

Xiang Liu, Simon W. Sanchez, Yan Gong, Roksana Riddle, Zebin Jiang, Stevens Trevor, Christopher H. Contag, Debajit Saha, and Wen Li. An insect-based bioelectronic sensing system combining flexible dual-sided microelectrode array and insect olfactory circuitry for human lung cancer detection. Biosensors and Bioelectronics, 281:117356, August 2025. ISSN 09565663. doi: 10.1016/j.bios.2025.117356. URL https://linkinghub.elsevier.com/retrieve/pii/S0956566325002301.

Valerio Mante, David Sussillo, Krishna V. Shenoy, and William T. Newsome. Context-dependent computation by recurrent dynamics in prefrontal cortex. Nature, 503(7474):78–84, November 2013. ISSN 0028-0836, 1476-4687. doi: 10.1038/nature12742. URL https://www.nature.com/articles/nature12742.

Ofer Mazor and Gilles Laurent. Transient dynamics versus fixed points in odor representations by locust antennal lobe projection neurons. Neuron, 48(4):661–673, 2005. ISSN 0896-6273. doi: 10.1016/j.neuron.2005.09.032. URL https://www.sciencedirect.com/science/article/pii/S0896627305008925.

Ofer Mazor, Gilles Laurent, and Gilles Laurent. Transient dynamics versus fixed points in odor representations by locust antennal lobe projection neurons. Neuron, 48(4):661–673, November 2005. doi: 10.1016/j.neuron.2005.09.032. MAG ID: 2030092068.

Summer B McLane-Svoboda, Shruti Joshi, Autumn K McLane-Svoboda, Camron Stout, Maksim Bazhenov, and Debajit Saha. Detection of multiple per-and polyfluoroalkyl substances (pfas) using a biological brain-based gas sensor. bioRxiv, pages 2025–09, 2025.

Katherine I. Nagel and Rachel I. Wilson. Mechanisms Underlying Population Response Dynamics in Inhibitory Interneurons of the Drosophila Antennal Lobe. The Journal of Neuroscience, 36(15):4325–4338, April 2016. ISSN 0270-6474, 1529-2401. doi: 10.1523/JNEUROSCI.3887-15.2016. URL https://www.jneurosci.org/lookup/doi/10.1523/JNEUROSCI.3887-15.2016.

Michael Parnas, Autumn K. McLane-Svoboda, Elyssa Cox, Summer B. McLane-Svoboda, Simon W. Sanchez, Alexander Farnum, Anthony Tundo, Nöel Lefevre, Sydney Miller, Emily Neeb, Christopher H. Contag, and Debajit Saha. Precision detection of select human lung cancer biomarkers and cell lines using honeybee olfactory neural circuitry as a novel gas sensor. Biosensors and Bioelectronics, 261:116466, October 2024. ISSN 09565663. doi: 10.1016/j.bios.2024.116466. URL https://linkinghub.elsevier.com/retrieve/pii/S0956566324004718.

Mainak Patel, Aaditya V. Rangan, and David Cai. A large-scale model of the locust antennal lobe. Journal of Computational Neuroscience, 27(3):553–567, December 2009. ISSN 0929-5313, 1573-6873. doi: 10.1007/s10827-009-0169-z. URL http://link.springer.com/10.1007/s10827-009-0169-z.

Matthew G. Perich, Charlotte Arlt, Sofia Soares, Megan E. Young, Clayton P. Mosher, Juri Minxha, Eugene Carter, Ueli Rutishauser, Peter H. Rudebeck, Christopher D. Harvey, and Kanaka Rajan. Inferring brain-wide interactions using data-constrained recurrent neural network models, December 2020. URL http://biorxiv.org/lookup/doi/10.1101/2020.12.18.423348.

Christophe Pouzat, Ofer Mazor, and Gilles Laurent. Using noise signature to optimize spike-sorting and to assess neuronal classification quality. Journal of Neuroscience Methods, 122(1):43–57, December 2002. ISSN 01650270. doi: 10.1016/S0165-0270(02)00276-5. URL https://linkinghub.elsevier.com/retrieve/pii/S0165027002002765.

Kanaka Rajan, Christopher D. Harvey, and David W. Tank. Recurrent Network Models of Sequence Generation and Memory. Neuron, 90(1):128–142, April 2016. ISSN 0896-6273. doi: 10.1016/j.neuron.2016.02.009. URL https://www.cell.com/neuron/abstract/S0896-6273(16)00102-1.

Debajit Saha, Kevin Leong, Chao Li, Steven Peterson, Gregory Siegel, and Baranidharan Raman. A spatiotemporal coding mechanism for background-invariant odor recognition. Nature neuroscience, 16(12): 1830–1839, 2013.

Debajit Saha, Wensheng Sun, Chao Li, Srinath Nizampatnam, William Padovano, Zhengdao Chen, Alex Chen, Ege Altan, Ray Lo, Dennis L. Barbour, and Baranidharan Raman. Engaging and disengaging recurrent inhibition coincides with sensing and unsensing of a sensory stimulus. Nature Communications, 8(1):15413, May 2017. ISSN 2041-1723. doi: 10.1038/ncomms15413. URL https://www.nature.com/articles/ncomms15413.

Simon W. Sanchez, Erin L. Vegter, Michael Parnas, Yong Song, Asgerally T. Fazleabas, and Debajit Saha. An insect brain-based bioelectronic neural sensor for the systemic detection and precise classification of endometriosis, June 2025. URL http://biorxiv.org/lookup/doi/10.1101/2025.06.03.657650.

Ben Scholl, Xiang Gao, and Michael Wehr. Nonoverlapping Sets of Synapses Drive On Responses and Off Responses in Auditory Cortex. Neuron, 65(3):412–421, February 2010. ISSN 08966273. doi: 10.1016/j.neuron.2010.01.020. URL https://linkinghub.elsevier.com/retrieve/pii/S0896627310000462.

Magdalena Solyga and Tania Rinaldi Barkat. Emergence and function of cortical offset responses in sound termination detection. eLife, 10:e72240, December 2021. ISSN 2050-084X. doi: 10.7554/eLife.72240. URL https://elifesciences.org/articles/72240.

David Sussillo and Omri Barak. Opening the Black Box: Low-Dimensional Dynamics in High-Dimensional Recurrent Neural Networks. Neural Computation, 25(3):626–649, March 2013. ISSN 0899-7667. doi: 10.1162/NECO_a_00409. URL https://doi.org/10.1162/NECO_a_00409.

David Sussillo, Mark M Churchland, Matthew T Kaufman, and Krishna V Shenoy. A neural network that finds a naturalistic solution for the production of muscle activity. Nature Neuroscience, 18(7):1025–1033, July 2015. ISSN 1097-6256, 1546-1726. doi: 10.1038/nn.4042. URL https://www.nature.com/articles/nn.4042.

Adrian Valente, Jonathan W Pillow, and Srdjan Ostojic. Extracting computational mechanisms from neural data using low-rank rnns. In S. Koyejo, S. Mohamed, A. Agarwal, D. Belgrave, K. Cho, and A. Oh, editors, Advances in Neural Information Processing Systems, volume 35, pages 24072–24086. Curran Associates, Inc., 2022. URL https://proceedings.neurips.cc/paper_files/paper/2022/file/9877d915a4b4f00e85e7b4cfdf41e450-Paper-Conference.pdf.

Daniel L. K. Yamins, Ha Hong, Charles F. Cadieu, Ethan A. Solomon, Darren Seibert, and James J. DiCarlo. Performance-optimized hierarchical models predict neural responses in higher visual cortex. Proceedings of the National Academy of Sciences, 111(23):8619–8624, June 2014. ISSN 0027-8424, 1091-6490. doi: 10.1073/pnas.1403112111. URL https://pnas.org/doi/full/10.1073/pnas.1403112111.

